# Single-Molecule Dynamics of SARS-CoV-2 5’ Cap Recognition by Human eIF4F

**DOI:** 10.1101/2021.05.26.445185

**Authors:** Hea Jin Hong, Matthew G. Guevara, Eric Lin, Seán E. O’Leary

## Abstract

Coronaviruses initiate translation through recognition of the viral RNA 5’ m^7^GpppA_m_ cap by translation factor eIF4F. eIF4F is a heterotrimeric protein complex with cap-binding, RNA-binding, and RNA helicase activities. Modulating eIF4F function through cellular regulation or small-molecule inhibition impacts coronavirus replication, including for SARS-CoV-2. Translation initiation involves highly coordinated dynamics of translation factors with messenger or viral RNA. However, how the eIF4F subunits coordinate on the initiation timescale to define cap-binding efficiency remains incompletely understood. Here we report that translation supported by the SARS-CoV-2 5’-UTR is highly sensitive to eIF4A inhibition by rocaglamide. Through a single-molecule fluorescence approach that reports on eIF4E–cap interaction, we dissect how eIF4F subunits contribute to cap-recognition efficiency on the SARS-CoV-2 5’ UTR. We find that free eIF4A enhances cap accessibility for eIF4E binding, but eIF4G alone does not change the kinetics of eIF4E–RNA interaction. Conversely, formation of the full eIF4F complex significantly alters eIF4E–cap interaction, suggesting that coordinated eIF4E and eIF4A activities establish the net eIF4F–cap recognition efficiency. Moreover, the eIF4F complex formed with phosphomimetic eIF4E(S209D) binds the viral UTR more efficiently than with wild-type eIF4E. These results highlight a dynamic interplay of eIF4F subunits and mRNA that determines cap-recognition efficiency.

## INTRODUCTION

A virus of unknown origin was first reported on December 31, 2019 in the city of Wuhan, China. Ensuing worldwide reports of the same virus alarmed the international community due to their rapid spread and associated sudden increase of severe pneumonia cases.^1^ The virus was confirmed to belong to the *Coronaviridae* family, and was initially named 2019-nCoV; it was later renamed as severe acute respiratory syndrome coronavirus 2 (SARS-CoV-2) and the associated disease was named COVID-19 by WHO.^1,2^ More than a year after the outbreak was declared a pandemic, more than 162 million confirmed cases have been reported, including more than 3.3 million deaths globally, in addition to significant impacts on the global economy.^2,3^

As obligate intracellular parasites, coronaviruses rely entirely on the host protein synthesis machinery for translation of viral proteins.^4^ The SARS-CoV-2 genome is a large, single-stranded positive-sense RNA – at 30 kb, one of the largest known viral RNA genomes.^5^ The positive-sense genomic RNA encodes, among others, 16 non-structural viral proteins.^5^ The viral RNA must compete with host mRNAs for cellular ribosomes in order to produce these proteins at the correct abundance and stoichiometry to ensure successful viral replication.^6^

Host-cell translation is also central to a major breakthrough in combating the SARS-CoV-2/COVID-19 pandemic: the rapid development of efficacious vaccines.^7^ Several vaccines with emergency approval for widespread use are mRNA-based.^8,9^ mRNA vaccines rely on host cell ribosomes and the associated translation apparatus to synthesize an antigenic viral protein fragment encoded by the vaccine mRNA in sufficient amounts to allow immune-system activation.^8^ Since in eukaryotes mRNA selection for translation is heavily regulated at its initiation step to control the efficiency of protein synthesis,^10^ understanding the molecular mechanisms of mRNA interaction with the human translation initiation machinery is important in understanding fundamental aspects of both the viral life cycle and vaccine efficacy.

Initiation on most cellular mRNAs involves recognition of an mRNA 5’ m^7^G(5’)ppp(5’)N cap structure^11,12^ where N is the +1 nucleotide of the transcript. The cap is recognized by the heterotrimeric eukaryotic translation initiation factor 4F (eIF4F).^13^ eIF4F binds the cap and recruits the small ribosomal subunit to mRNA, forming a 48S pre-initiation complex which is thought to scan toward and locate the start codon for ribosomal subunit joining and translation elongation.^14–16^ eIF4F is composed of a cap-binding subunit, eIF4E, which interacts with the terminal *N*^7^-methylguanosine base and the triphosphate bridge.^17–20^ eIF4E also binds the large, multidomain eIF4F “scaffold” subunit, eIF4G, which also possesses domains that bind eIF4A, eIF3, the poly(A)-binding protein, and RNA;^21–24^ in humans the eIF4G interaction with 40S subunit-bound eIF3 leads to physical attachment of the pre-initiation complex to the mRNA.^14,25,26^ The third eIF4F subunit, eIF4A, is a DEAD-box RNA helicase.^13,27,28^

mRNAs encoding different proteins vary in their dependence on eIF4F for translation efficiency – in particular, greater extent of structure in the 5’-untranslated region is associated with a greater relative eIF4F requirement for efficient translation.^13,29^ Physical accessibility of the 5’ cap due to varying structural propensity has been proposed to enhance the recruitment efficiency of translation factors to the mRNA, and thus of translation initiation.^13,14,30^ mRNA sequence elements have also been proposed to modulate cap-recognition efficiency.^31–33^ However, the molecular logic relating specific mRNA features to cap-recognition efficiency remains incompletely understood.

Owing to their centrality in the initiation mechanism, eIF4F subunits are important components of translational control mechanisms. Both cell-signaling pathways and small-molecule inhibitors modulate eIF4E activity to rapidly alter cellular translation patterns. eIF4E is phosphorylated at Ser^209^, in an eIF4G-dependent manner, by MNK (MAP kinase-interacting) kinases MNK1 and MNK2, and downstream of the p38 MAP kinase and/or ERK cell signaling pathway.^34^ Phospho-eIF4E has been found associated with polysomes and has been implicated in enhancing translation,^35^ though the specific role of eIF4E phosphorylation in promoting translation has been questioned.^36^ Nevertheless, small-molecule eIF4F inhibitors have been of significant interest, particularly as anticancer agents.^37^

Coronaviruses, including SARS-CoV-1, initiate translation through a cap- and eIF4F-dependent mechanism.^38^ Accordingly, disruption of eIF4E–eIF4G interaction by the small-molecule inhibitor 4E2RCat abolished HCoV-229E replication in a cell-based assay, while host protein synthesis was inhibited only by 40%.^39^ Likewise, the eIF4A inhibitor silvestrol inhibited expression of MERS-CoV and HCoV-229E nonstructural proteins and formation of viral transcription complexes.^40^ eIF4F inhibition was also identified in a comprehensive survey of potential small-molecule therapeutic strategies for SARS-CoV-2.^6^ However, the dynamics of eIF4F interaction with a capped coronavirus 5’-untranslated region have not been elucidated.

Coronavirus infection activates the p38 pathway MAP kinase, leading to eIF4E phosphorylation at residue Ser^209^.^38^ However, while eIF4E phosphorylation is frequently thought to promote translation,^41^ global cellular protein synthesis is attenuated following coronavirus infection.^38^ Furthermore, a MAP kinase inhibitor failed to reduce viral translation in SARS-CoV-1-infected cells even though it inhibited MAP kinase-dependent eIF4E phosphorylation.^42^ Thus, the role of eIF4E phosphorylation specifically in coronavirus translation remains unclear.

In this study we developed a single-molecule fluorescence assay for human eIF4E interaction with capped mRNA, to gain insights into how the eIF4F subunits coordinate in cap recognition. We then applied this assay to characterize eIF4F interaction with the SARS-CoV-2 5’ untranslated region.

## MATERIAL AND METHODS

### Cloning and synthesis of vRNA constructs

DNA templates for *in vitro* RNA transcription of SARS-CoV-2 RNAs were obtained commercially (Genewiz, Inc.). For preparation of the luciferase fusion construct, DNA fragments containing the class II T7 promoter Φ2.5, followed by the SARS-CoV-2 5’ untranslated region sequence, the coding sequence for firefly luciferase, and the SARS-CoV-2 5’ untranslated region sequence were assembled in a pUC57 vector. A HindIII restriction site was placed at the 3’ end of the insert, to allow template linearization prior to transcription. The construct for transcription of the isolated 5’ UTR was prepared by the same strategy, by inserting class II T7 promoter Φ2.5 at the 5’ end and an EcoRI restriction site at the 3’ end to linearize the DNA template.

The full-length human *GAPDH* transcript sequence, obtained from UCSC Genome Browser, was purchased as a gBlock from IDT. The sequence was inserted into pUC119 *via* SalI and EcoRI restriction sites incorportated during gBlock synthesis. A T7 promoter sequence was also included at the 5’ end, along with an extra 5’ dG to facilitate efficient *in vitro* transcription. The EcoRI site placed at the 3’ end allowed linearization of the template for the subsequent transcription reaction.

Plasmids were isolated at preparative scale from *E*. coli DH5α transformants selected with ampicillin (100 µg/mL), using a maxi-prep kit (Macherey Nagel). The insert DNA sequences for each construct are given in Supplementary Information. The sequences of all plasmid inserts used in this study were confirmed by Sanger sequencing (Genewiz, Inc.).

### RNA transcription

Purified plasmids were digested at the 3’ end of the insert sequence with EcoRI-HF (NEB) to linearize the template with a 5’ overhang for in vitro transcription. For 60 µL transcription reactions, about 11 µg of linearized DNA template was incubated for 4 h at 37 °C with 12.5 mM of each NTP, 3% (v/v) DMSO, 25 mM MgCl_2_, 17 mM DTT, and T7 RNA polymerase (12,500 units/mL) in transcription buffer composed of 0.4 M Tris-HCl (pH 8.1), 10 mM spermidine, and 0.01% (v/v) Triton X-100. The transcription product was extracted using acidic phenol chloroform (pH 4.5) and precipitated overnight in ethanol at –20 °C. RNA was redissolved in water. RNAs were capped using the *Vaccinia* capping system (NEB) and poly(A)-tailed at their 3’ ends with poly(A) polymerase (NEB), following the manufacturer’s protocol. RNA concentration was quantified by UV absorbance spectrophotometry using a NanoDrop instrument.

### Luciferase assays

100 nM CoV firefly luciferase fusion RNA was added to 12 µL of HeLa lysate reaction mix from an *in vitro* protein expression kit (Thermo Scientific, 88882), along with 0.5 µL of rocaglamide/RocA solution (Med Chem Express, Product No. HY-19356) or 4E2RCat solution (Med Chem Express, Product No. HY-100733) in varying concentrations as indicated in the Results and Discussion. RocA and 4E2RCat stock solutions were prepared in 10% (v/v) and 100% DMSO, respectively; the final DMSO concentration in all reactions was 0.4% and 4 % respectively (v/v). The reaction was incubated at 30 °C for 5 hours and 2.5 µL of the reaction mix was added to 50 µL of luciferase assay reagent (Promega) pre-equilibrated in room temperature. The luminescence was then measured using a Luminometer Turner (Turner Biosystems). Luciferase assays were performed in triplicate and the luminescence of each replicate was determined as the average of three separate measurements. Luminescence was normalized to a DMSO-only control reaction, by dividing the luminescence measured from the reactions containing the inhibitors with the luminescence measured from the sample only containing the DMSO at the appropriate concentration for each inhibitor.

### Preparation of fluorescently labeled eIF4E and eIF4E(S209D)

The pET-28a(+) vectors containing the sequences of eIF4E or eIF4E(S209D), each fused with an N-terminal Protein G tag, were designed similarly to Feokistova *et. al*., (2013).^28^ The constructs contained a sequence encoding the peptide MA(*p*AzF) between the Protein G tag and the N-terminus of eIF4E, where *p*AzF is *p*-azidophenylalanine (pAzF). *p*AzF was encoded by the amber stop codon (TAG), for decoding by *p*AzF-tRNA^CUA^. The insert sequences are given in the Supplementary Information.

The pULTRA expression vector, containing inserts encoding tRNA^CUA^ and pAzF-tRNA^CUA^-synthetase – a generous gift from Abhishek Chatterjee, Boston College – was co-transformed into an *E. coli* BL95ΔAΔ*fabR* strain with the pET28a(+)-eIF4E plasmid, and transformants were selected on LB agar containing Kanamycin (50 µg/mL) and Spectinomycin (100 µg/mL). A single colony from this selection was used to inoculate a 10 mL 2×YT broth starter culture containing both antibiotics, which was grown overnight at 37 °C. The following day, the starter culture was used to inoculate 1 L of 2xYT broth containing the antibiotics. The culture was grown at 37 °C to OD_600_ ∼0.6, and then 1 mM pAzF (BAChem, F3075) was added along with 1 mM isopropyl-β-D-thiogalactopyranoside (IPTG) to induce recombinant protein overexpression. Induction was carried out for 5 h at 30 °C in darkness. The cells were spun down and suspended in lysis buffer (20 mM HEPES-KOH pH 7.5, 400 mM KCl, 5 mM imidazole, 1×EDTA-freeprotease inhibitor cocktail (0.24 mg/mL benzamidine hydrochloride hydrate, 2 µM pepstatin, and 0.6 µM leupeptin hemisulfate), 1× phenylmethylsulfonyl fluoride (PMSF)). Proteins were purified immediately after induction, without freezing of the cell pellet. Further purification was carried out on ice with ice-chilled buffers. The cells were lysed using a sonicator equipped with a microtip (Branson Sonifier 450; 80 % amplitude, setting 3, for 45 seconds at 5 min intervals) and the lysate was immediately clarified by centrifugation (22,000 rpm for 30 min). The filtered lysate was passed through 5 mL Ni-NTA agarose (Thermo Fisher), followed by typically 3 washes with 5 column volumes of wash buffer (20 mM HEPES-KOH pH 7.5, 400 mM KCl, 10 mM imidazole) until no further detectable protein eluted from the column, as assessed by Bradford reagent (Bio-Rad). Ni-NTA-bound protein was then eluted with elution buffer (20 mM HEPES-KOH pH 7.5, 400 mM KCl, 200 mM imidazole), and eluate fractions were analysed by 10 % SDS-PAGE. Fractions that contained eIF4E were combined, and labeled overnight at 4 °C with 50 µM Cy5-DBCO (Sigma). The fluorophore was dissolved in 100% DMSO for addition to the labelling reaction; the final DMSO concentration in the labelling reaction was 1% (v/v). The labeled protein was desalted to remove unreacted dye and the sample was buffer exchanged into TEV cleavage buffer (20 mM HEPES-KOH, 50 mM KCl, 10 % (v/v) glycerol). The desalted sample was then incubated with TEV protease (NEB) (50 units) overnight at 4 °C for complete cleavage of protein G from the 4E. Once the cleavage was complete, the sample was loaded onto an SP-column (GE Healthcare Life Sciences) maintained at ∼4 °C. The column was washed with 20 column volumes of low-salt buffer (20 mM HEPES-KOH, 50 mM KCl, 10 % Glycerol), then eluted with a linear gradient from 0% to 100% high-salt buffer (20 mM HEPES-KOH, 500 mM KCl, 10 % Glycerol). Fractions containing Cy5-eIF4E typically eluted at ∼250 mM KCl. Based on inspection of an SDS-PAGE gel of the eluate fractions, imaged for Cy5 fluorescence by a Typhoon imager, Fractions containing labeled protein were combined, concentrated to < 1 mL by centrifugal ultrafiltration (10,000 or 5,000 MWCO ultrafilter) and loaded onto a Superdex 75 Increase (10/300 GL) column (GE Healthcare Life Sciences) equilibrated in storage buffer (20 mM HEPES-KOH pH 7.5, 200 mM KCl, 10 % Glycerol, 1 mM TCEP) for further purification. Fractions were assessed for purity by SDS-PAGE, and for labelling efficiency by measuring the ratio of 640 nm to 280 nm absorbance. The 1,2,3-triazole formed in the azide-alkyne cycloaddition reaction that conjugates the fluorophore to the protein is expected to absorb at 280 nm. Considering the contribution of this additional absorbance, the typical labelling efficiency was ∼50%; this assessment was supported by SDS-PAGE analysis of the labelled protein, which contained two bands, only one of which was conjugated to Cy5 (Fig. S1A). Purified Cy5-eIF4E and Cy5-eIF4E(S209D) were stored at 4 °C in darkness and used for a maximum of one week after each purification.

### Preparation of human eIF4A

pHis APEX2-eIF4A1 was a gift from Nicholas Ingolia (Addgene plasmid #12964; Ingolia et. al., 2019).^43^ The plasmid was transformed into BL21(DE3) CodonPlus cells and transformants were selected on LB–agar plates containing ampicillin (100 µg/mL) and chloramphenicol (25 µg/mL). A single colony from this selection was used to grow a 10 mL overnight culture at 37 °C. This starter culture was then used to inoculate 1 L LB containing the selective antibiotics. The cells were grown to OD_600_ of ∼0.6, then protein overexpression was induced with 1 mM IPTG for 3 h at 37 °C. After induction, the cells were harvested by centrifugation at 4,000 rpm for 15 min and were frozen and stored at -80 °C until purification. The frozen cell pellet was resuspended in lysis buffer (20 mM HEPES-KOH, pH 7.5, 500 mM NaCl, 10 mM imidazole, 10 mM β-mercaptoethanol, 0.5% (v/v) NP-40). The cells were lysed by sonication with a microtip attachment (80 % amplitude, setting 3, for 45 seconds at 5 min intervals) and the lysate was immediately clarified by centrifugation (22,000 rpm for 30 min). During centrifugation, a gravity-flow column containing 1 mL Ni-NTA agarose (Thermo Scientific) was washed and pre-equilibrated with the lysis buffer. The cell-free supernatant was diluted twofold with this lysis buffer, and filtered through a 0.22 µm syringe filter (Corning). The filtrate was then applied to the equilibrated Ni-NTA column, which was washed with a further ∼20 mL lysis buffer. The column was further washed with 40 mL high-salt buffer (20 mM HEPES-KOH, pH 7.5, 1 M NaCl, 20 mM imidazole, 10 mM β-mercaptoethanol), followed by low-salt buffer (20 mM HEPES-KOH, pH 7.5, 500 mM NaCl, 20 mM imidazole, 10 mM β-mercaptoethanol). The protein was then eluted with elution buffer (50 mM Na-phosphate buffer, pH 7.5, 500 mM NaCl, 100 mM Na2SO4, 250 mM imidazole, 2 mM DTT). After elution, the eluate was buffer-exchanged into low-salt buffer (20 mM Tris HCl pH 7.5, 100 mM KCl, 5% glycerol, 2 mM DTT, 0.1 mM EDTA) using a 10 DG column (Bio-Rad), and was TEV-protease (50 units; NEB) cleaved overnight at 4°C to remove the APEX tag. The TEV-protease cleaved sample was loaded onto a 5 mL Q-Sepharose HP column (GE Healthcare Life Sciences), equilibrated in low-salt buffer and maintained at ∼4 °C. The column was washed with 20 column volumes of low-salt buffer, then eluted with a linear gradient from 0% to 100% high-salt buffer (20 mM Tris HCl pH 7.5, 500 mM KCl, 5% glycerol, 2 mM DTT, 0.1 mM EDTA) over 50 column volumes. eIF4A typically eluted at ∼265 mM KCl. Eluate fractions were analyzed by SDS-PAGE, and fractions containing pure eIF4A were then buffer exchanged into a storage buffer (20 mM Tris HCl, pH 7.5, 2 mM DTT, 0.1 mM EDTA, 10% glycerol, 100 mM KCl) using Superdex-75, flash frozen under liquid nitrogen and was stored at -80 °C.

### Preparation of His_6_-tagged human eIF4G(557-1137)

A pET-28a(+) vector was constructed with an insert containing a sequence encoding a hexahistidine tag, followed by the sequence encoding residues 557 to 1137 of human eIF4G1, codon-optimized for *E. coli*.. The plasmid was transformed into BL21 (DE3) CodonPlus cells. Transformants were selected on LB-agar plates containing kanamycin (50 µg/mL) and chloramphenicol (25 µg/mL). A single transformant colony was used to inoculate a 10 mL LB starter culture, which was grown overnight at 37 °C. The next day the starter culture was used to inoculate six 1 L LB cultures containing the selective antibiotics. Cells were grown at 37°C to OD_600_ of ∼0.6, then protein overexpression was induced by addition of 1 mM IPTG, and allowed to proceed overnight at 16 °C. The cells were pelleted by centrifugation at 4,000 rpm for 15 minutes and stored at -80°C until purification. Frozen cell pellets were resuspended in lysis buffer (20 mM sodium phosphate, pH 7.5, 300 mM NaCl, 10% (v/v) glycerol, 10mM β-mercaptoethanol, 1× EDTA-free protease inhibitor (Roche), 1× PMSF, 10mM imidazole) and lysed using the microtip sonicator described above (80 % amplitude, setting 3, for 30 seconds in 2 minute intervals). Approximately 3 mL Ni-NTA agarose (Thermo Scientific) was washed and pre-equilibrated with lysis buffer while the lysate was clarified by centrifugation (22,000 rpm for 30 min). The clarified lysate was filtered first through 0.80 µm syringe filter (Corning) then by a 0.22 µm filter. The filtered lysate was added to the equilibrated column and washed with ∼15 mL lysis buffer. At room temperature, the column was pre-treated with 25 mL micrococcal nuclease buffer (20 mM sodium phosphate, pH 7.5, 100 mM NaCl, 2 mM CaCl_2_) prior to treating with 1.5 mL micrococcal nuclease buffer containing 3 U/µL micrococcal nuclease (pre-warmed to 37°C) followed by immediate incubation at 37°C for 30 mins. After this incubation, at room temperature, the column was washed with lysis buffer supplemented with 5 mM EGTA to stop the reaction. The remaining washes (15 mL each) were carried out at 4 °C: wash buffer 1 (20 mM sodium phosphate, pH 7.5, 800 mM NaCl, 10% (v/v) glycerol, 10 mM β-mercaptoethanol), wash buffer 2 (20 mM sodium phosphate, pH 7.5, 100 mM NaCl, 10 % Glycerol, β-mercaptoethanol), wash buffer 3 (20 mM sodium phosphate, pH 7.5, 100 mM NaCl, 10% (v/v) glycerol, 20 mM imidazole, 10 mM β-mercaptoethanol). Ni-NTA-bound proteins were eluted with 0.75 mL aliquots of elution buffer (20 mM sodium phosphate, pH 7.5, 100 mM NaCl, 10% (v/v) glycerol, 500 mM Imidazole) and DTT was added to each elution fraction to a final concentration of 2 mM. Protein fractions containing the eIF4G fragment, as assessed by SDS-PAGE, were pooled and buffer exchanged to remove excess imidazole (20mM sodium phosphate, pH 7.5, 100mM NaCl, 10% glycerol) using a 10-DG column (Bio-Rad; pre-equilibrated in elution buffer without imidazole and DTT). The protein eluted from the 10-DG column (3.5 mL) was divided into four samples, which were each applied to ∼400 µL of magnetic Ni-NTA beads (NEB, product number S1423S), equilibrated at 4 °C in binding buffer (50 mM sodium phosphate, 300 mL NaCl, 10 mM imidazole, pH 8.0). The mixture was rocked on ice for one hour, then the supernatant was removed. The beads were washed with three 500 µL aliquots of wash buffer (50 mM sodium phosphate, 300 mM NaCl, 20 mM imidazole, pH 8.0), then bound proteins were eluted by rocking the beads on ice for 10 minutes after addition of 100 µL of elution buffer (50 mM sodium phosphate, 300 mM NaCl, 500 mM imidazole, pH 8.0). This magnetic-bead step allowed the protein to be concentrated under mild conditions; repeated attempts at concentration by centrifugal ultrafiltration resulted in very low levels of protein recovery. The resulting protein was then dialyzed (Thermo Scientific; Product No. 69552) into storage buffer (20 mM sodium phosphate pH 7.5, 100 mM NaCl, 10 % glycerol, 2 mM DTT), flash-frozen in liquid nitrogen, and stored at –80 °C until use.

### Electrophoretic mobility shift assay

25 nM SARS-CoV-2 5’ UTR was added to varying concentration of eIF4G(557-1137) (0 to 300 nM) in 1 x binding buffer (40 mM HEPES-KOH pH 7.5, 25 mM KCl, 1 mM EDTA, 2.5 mM MgCl_2_, 0.3 mM DTT, 0.01% (v/v) NP-40). Additional salt was added using eIF4G storage buffer, to bring the KCl concentration to100 mM, and glycerol concentration to 10 % (v/v) in all reactions, with a total volume of 15 µL. The binding reaction was incubated in a 30 °C incubator for 8 min. The samples were then loaded onto a 4% polyacrylamide minigel (8.6 × 6.7 × 0.75 cm) (19:1 acrylamide:bisacrylamide) prepared with THEM buffer (34 mM Tris base, 57 mM HEPES, 0.1 mM EDTA, 2.5 mM Mg(OAc)_2_, pH 7.5). The gel was pre-run at 80V for 30 minutes, at 4 °C, then the binding reaction mixtures were loaded and the gel was electrophoresed at 80 V for 1 h at 4°C. The gel was post-stained with 0.5 x Gel-Red (Biotium, 41001) for a few seconds and rapidly visualized on a Bio-Rad ChemiDoc gel imager, using the EtBr filter.

### eIF4A ATPase assay

An NADH-linked enzyme coupled assay^44^ was performed to measure the ATPase activity of eIF4A. All reagents were freshly prepared, including the stocks of 15 mM NADH, 100 mM Mg•ATP (pH 7.0), Lactate dehydrogenase (LDH; 4000 U/mL; Sigma 427217.), 100 mM phosphoenolpyruvate (PEP; pH 7.0) and 1M DTT. A 5 x coupling assay cocktail was prepared in 1 x KMg75 buffer (20 mM HEPES-KOH pH 7.5, 75 mM KCl, 5 mM MgCl_2_ and 1 mM DTT), with the following components in final concentrations: 1 mM NADH, 100 U/mL LDH, 500 U/mL pyruvate kinase, 2.5 mM PEP. A separate tube containing 2 x coupling assay cocktail with or without proteins (eIF4A or eIF4G or both) in 1 x KMg75 buffer and another tube containing 2 mM ATP and 2 µM CoV RNA in 1 x KMg75 buffer were prepared. The solutions were mixed in a 100 µL quartz cuvette with 1 cm pathlength and immediately monitored using a UV spectrophotometer (Shimadzu) at 340 nm for 15 min, at 0.7-s time intervals. The time course of absorbance change was then fit by linear regression to obtain the slope. This and the background NADH conversion (i.e., in the absence of proteins) were used to compute the observed ATPase rates with the following formulae:

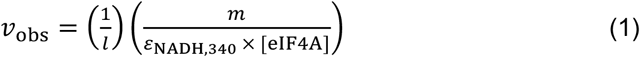

*l* is the pathlength, *m* is the background corrected slope, *ε*_NADH.340_ is the molar absorptivity of NADH at 340 nm, (6,220 mM^−1^ cm^−1^) and [eIF4A] is the total concentration of eIF4A added in the experiment. The reported values of *V* / *E*_o_ represent the average of three independent replicate experiments; error bars reflect the standard deviation of these replicates.

### Single-molecule experiments

A customized Pacific Biosciences RS II instrument was used to monitor real-time eIF4E interaction with the 5’ UTR of the viral RNA.^45–47^ For the experiment, a zero-mode waveguide (ZMW) chip (Pacific Biosciences SMRT Cell; part number 100-171-800) was first treated with NeutrAvidin mixture (16.6 µM neutravidin, 0.67 mg/mL BSA) in smBuffer 30 mM HEPES-KOH pH 7.5, 3 mM Mg(OAc)_2_ and 100 mM KOAc) at room temperature for 5 min, which allowed the NeutrAvidin to bind the biotin-PEG layer on base of the chip. Polyadenylated viral RNA was prepared for immobilization through annealing to biotin-5’-(dT)_45_-3’-Cy3 (purchased from IDT), by heating a mixture containing 1 µM RNA and 100 nM oligo in 0.1 M BisTris pH 7.0, 0.3 M KCl to 98 °C for 2 minutes in a thermocycler, then slow-cooling to 4 °C over 20 minutes. The resulting hybrid duplex was immobilized by incubating 20 µL of the annealing mixture (10 nM RNA mixture final) on the chip surface for ∼10 min at room temperature. The immobilization mixture was then removed, and the chip was washed three times with smBuffer. The chip was then pre-blocked with 1 µM unlabeled eIF4E, 5% (v/v) each Biolipidure 203 and 206 (NOF America Corporation), and 5 mg/mL purified BSA (NEB, G9001S), to prevent non-specific interaction of Cy5-eIF4E with the chip surface. After pre-blocking, the chip was washed a further three times with smBuffer. After the third wash, 20 µL was added to the chip surface of smBuffer supplemented with BSA, with an oxygen scavenging system (2.5 mM PCA (protocatechuic acid), 1× PCD (protocatechuate-3,4-dioxygenase, Pacific Biosciences)), and with triplet-state quencher (2 mM TSY, Pacific Biosciences). After loading the chip onto the instrument, the instrument delivered 60 nM Cy5 labeled hs4E WT or S209D mutant to the chip, along with unlabelled components as needed (i.e., eIF4G, eIF4A, and RocA), in a volume of 20 µL (final concentrations in the experiment are half the delivery concentrations, e.g. 30 nM eIF4E, due to dilution into the volume already on the chip surface). Where included, eIF4G(557-1137) and eIF4A were present at final concentrations of 40 nM and 1 µM, respectively. Movies were acquired for 10 minutes at 10 frames per second, under illumination with a 532 nm laser at a power of 0.70 µW/µm^2^, to excite Cy3 fluorophores. In the experiment with direct red illumination (Figure S3), the 642 nm laser power was 0.07 µW/µm^2^.

### Single-molecule data analysis

Single-molecule fluorescence traces were extracted from raw movie files with custom MATLAB scripts reported previously.^45^ Traces containing robust smFRET signals for eIF4E–RNA interaction were selected by visual inspection. Selection criteria were: a stable Cy3 signal at the very beginning of the movie, a single Cy3 photobleaching event, and visually apparent FRET to Cy5. Assignment of event timings in the traces was carried out using a hidden Markov model approach as implemented in ebFRET.^48^ Event timings identified from ebFRET analysis were converted to empirical cumulative probability distributions for the times between events, and the event durations, then fit with single- or double-exponential models to determine *k*_on_ or *k*_off_, according to the equations:

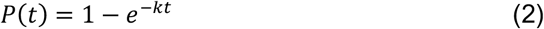

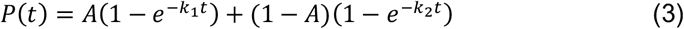

Fitting was carried out in MATLAB by non-linear least-squares regression using the Trust-Region algorithm. Goodness-of-fit was confirmed by inspection of the root-mean-squared errors for the fits, which were 0.01 – 0.03 in all cases. Values for fitted rate constants in each experimental condition are given in Supplementary Table 1.

## RESULTS AND DISCUSSION

### SARS-CoV-2 untranslated regions support eIF4F-dependent translation

To formally confirm that the SARS-CoV-2 untranslated regions supports eIF4F-dependent translation in our hands, in analogy to other coronaviruses including SARS-CoV-1, we designed a reporter mRNA containing firefly luciferase bracketed by the viral 5’ and 3’ UTR regions as annotated in the SARS-CoV-2 reference genome (Fig. 1A).^49^ The mRNA was prepared with the m^7^GpppA_m_ cap structure (cap-1) found on coronavirus RNAs in cells.^50^ The luciferase construct contained a 33-nucleotide poly(A) tail encoded by the DNA transcription template.

**Figure 1.**
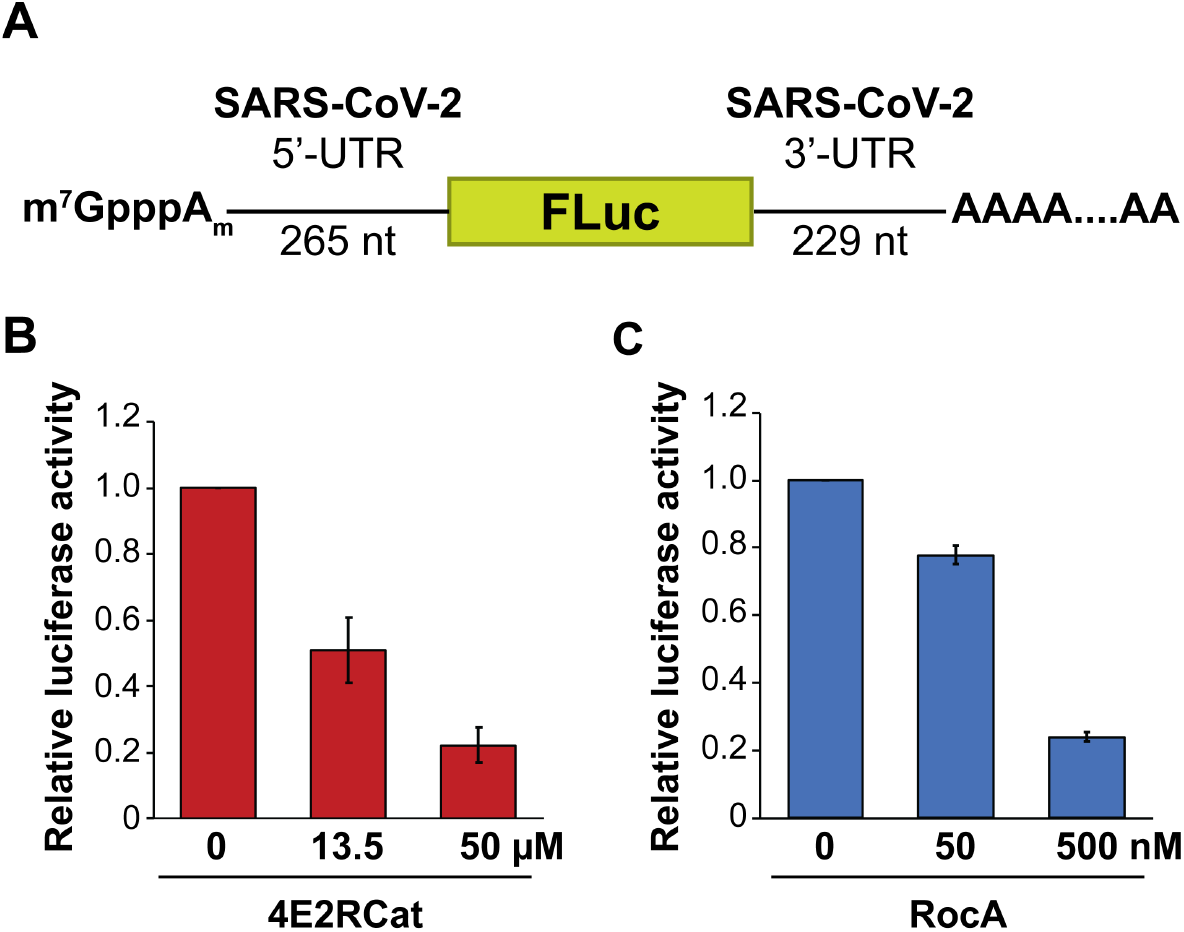
SARS-CoV-2 untranslated regions support eIF4F-dependent translation. (A) Schematic of luciferase reporter construct. (B) Inhibition of reporter translation in a HeLa cell-free translation assay by the small-molecule inhibitor of eIF4E–eIF4G interaction, 4E2RCat. Luciferase translation is normalized to a control reaction without inhibitor. (C) Inhibition of reporter translation by the small-molecule eIF4A inhibitor, RocA (rocaglamide). Error bars in (B) and (C) reflect the standard deviation from three independent luciferase reactions, with the luminescence from each independent reaction measured in triplicate.

We assayed the effects of the small-molecule inhibitors 4E2RCat and RocA on translation of the reporter mRNA in a HeLa cell-free translation system. 4E2RCat inhibits eIF4E–eIF4G interaction with an IC_50_ value of 13.5 µM, and inhibits HCoV-229E replication.^39^ RocA, or rocaglamide, is a member of the rocaglate/flavagline class of eIF4A inhibitors, produced in *Aglaia* species, which lock the enzyme onto polypurine mRNA sequences.^51^ This enhanced RNA affinity is thought to convert eIF4A into a physical block to ribosomal scanning.^52,53^ Rocaglates have also been proposed to sequester eIF4F on the mRNA cap.^54^

Both 4E2RCat and RocA abrogated reporter translation driven by the SARS-CoV-2 untranslated regions. Translation was reduced by ∼50% at 13.5 µM 4E2RCat (Fig. 1B), consistent with the previous studies. However, RocA inhibited translation much more potently, with a ∼25% reduction at 50 nM RocA (Fig. 1C). Thus, the viral untranslated regions support eIF4F-dependent translation. Human mRNAs show widely varying sensitivity to RocA translation inhibition, with IC_50_ values ranging from ∼3 nM to 50 nM.^53,55^ Our data, although obtained with the isolated viral UTRs decoupled from the context of the full viral genome, suggest SARS-CoV-2 translation is similarly sensitive to RocA inhibition, echoing identification of the related rocaglate, silvestrol, as a potential SARS-CoV-2 therapeutic,^6^ the finding that the synthetic rocaglate CR-31-B(–) inhibits SARS-CoV-2 translation,^56^ the potent inhibitory activity of RocA against other coronaviruses,^40^ and the ability of rocaglates to selectively target pathologies resulting from aberrant translation.^52^

### A single-molecule fluorescence approach to observe human eIF4F-vRNA interaction

We next developed an assay to observe real-time human eIF4E/eIF4F recognition of the viral 5’-UTR, extending our previous single-molecule fluorescence approach for yeast eIF4E.^46,47^ In this assay (Fig. 2A), the capped SARS-CoV-2 5’-UTR is polyadenylated at its 3’ end, then hybridized to biotin-5’-(dT)_45_-3’-Cy3, which both fluorescently labels it (Cy3) and allows it to be surface-immobilized through biotin-avidin interactions.^47^ Cy3 also serves as a donor fluorophore for Förster Resonance Energy Transfer (FRET). Real-time Cy5-eIF4E binding to single Cy3-mRNA molecules is observed through appearance of eIF4E–RNA FRET (Fig. 2B) when the Cy5-labeled factor binds the cap structure of the Cy3-labeled RNA, allowing association and dissociation rates to be determined. The assay is performed in zero-mode waveguides (ZMWs) using a customized Pacific Bioscience RS II instrument.^45^

**Figure 2.**
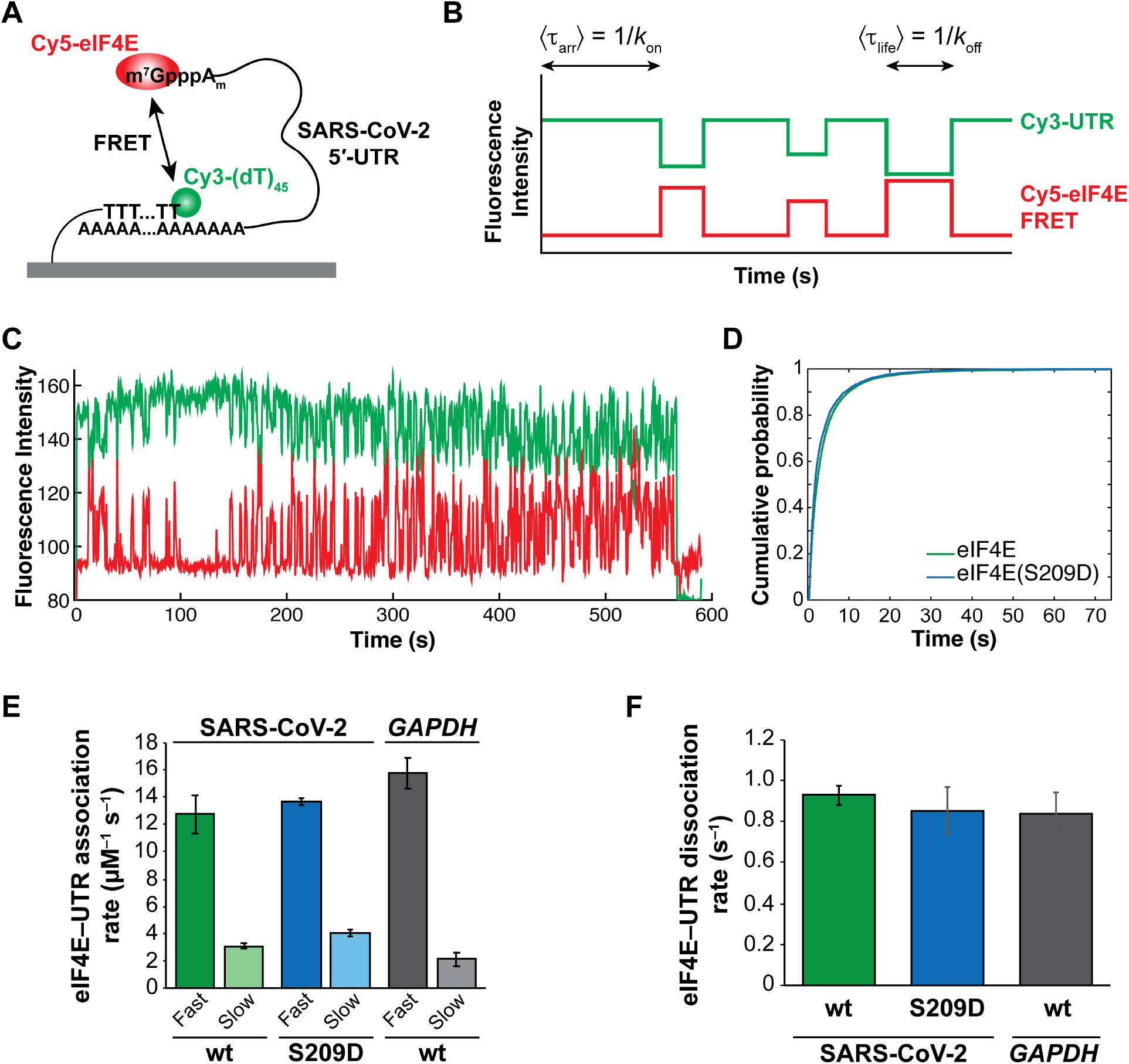
Single-molecule dynamics of human eIF4E interaction with the SARS-CoV-2 5’-UTR. (A) Schematic of experimental design, showing immobilization strategy and the basis of the smFRET signal for eIF4E–UTR binding. (B) Schematic idealized single-molecule fluorescence trace, showing the FRET signal obtained upon eIF4E–UTR binding. The mean waiting time between FRET events provides the eIF4E association rate. The mean duration of FRET events provides the dissociation rate. (C) Representative single-molecule fluorescence trace for eIF4E interaction with the surface-immobilized viral UTR. (D) Cumulative distribution function for arrival times of eIF4E (green) and eIF4E(S209D) (blue) to the surface-immobilized viral UTR. (E) Second-order rate constants calculated for association of eIF4E and eIF4E(S209D) with the viral UTR, and for eIF4E with full-length human *GAPDH* mRNA. Observed rates were determined by fitting of eIF4E-UTR arrival-time distributions to a double-exponential model by nonlinear least-squares regression. (F) Rates of eIF4E and eIF4E(S209D) dissociation from the viral UTR and *GAPDH*. Error bars in (D) and (E), and kinetic data throughout, reflect standard deviations from analysis of at least two independent experiments. Numbers of molecules analysed for each replicate are given in Supplementary Table 1.

We generated Cy5-labeled human eIF4E (Fig. S1A) by incorporation of a *p*-azidophenylalanine amino acid residue at the protein’s N terminus, through co-expression of an orthogonal tRNA/aminoacyl-tRNA transferase pair during eIF4E overexpression in *E. coli*.^57,58^ The azide-bearing protein was then labeled under mild conditions during purification, in a strain-promoted click reaction with a Cy5-DBCO alkyne. The resulting protein contains the native eIF4E sequence, extended at its N-terminus by the nonapeptide GHMA(*pAz*)FMLE. In addition, to gain insights into the role of eIF4E Ser^209^ phosphorylation, we prepared the phosphomimetic variant eIF4E(S209D) which we fluorescently labeled with the same approach (Fig. S1A). eIF4E(S209D) overexpression recapitulates cellular phenotypes attributable to eIF4E phosphorylation in a range of contexts,^59–61^ and the serine-to-aspartate substitution reduced the affinity of eIF4E for a capped RNA oligonucleotide.^62^ We thus reasoned that this protein was a suitable proxy for phospho-eIF4E in our assay. Both proteins were isolated in a monomeric form after size-exclusion chromatography (Fig. S1B,C); this is important as eIF4E aggregation modulates its interaction with the cap structure.^63^

### eIF4E dynamically interacts with SARS-CoV-2 RNA

Cy5-eIF4E interaction with surface-immobilized viral RNA was characterized by transient eIF4E binding to and dissociation from the cap (Fig. 2C), and resembling the interaction for the yeast protein.^46^ We previously demonstrated that RNA 5’ and 3’ ends remain situated within FRET distance (2 – 8 nm) in this assay; thus, the FRET signal does not significantly “miss” binding events due to intervals where the mRNA ends have moved out of FRET range.^47^ Experiments detailed below with the full eIF4F complex also did not show evidence for a significant extent of eIF4E–cap binding without FRET. While we cannot rule out that extremely transient events are not observed, our kinetic data are consistent overall with the affinity of eIF4E for capped RNAs,^63–65^ further strongly suggesting that our signal accurately recapitulates the interaction dynamics.

The distribution of times between eIF4E–UTR FRET events for many cycles of binding over ≥ 99 UTR molecules showed two populations (Fig. 2D). We calculated the association rates by fitting the arrival-time distribution to a double-exponential model; the fitted mean of each exponential component corresponds to the inverse of its observed rate constant, i.e. ⟨τ⟩ = 1/*k*. The relative incidence of events occurring at different rates is represented by the amplitude of the corresponding exponential phase in the arrival-time distribution, provided the acquisition time is sufficiently long to allow robust sampling of each rate process. At the eIF4E concentrations employed in the present study, the acquisition time was almost two orders of magnitude longer than the time constants ⟨τ⟩ for the slow-binding rate process.

Most (∼82 ± 6%) of the eIF4E–UTR binding events occurred at a rate of 12.7 ± 1.4 µM^−1^ s^−1^, whilst the remainder occurred at 3.1 ± 0.2 µM^−1^ s^−1^ (Fig. 2E; Supplementary Table 1). The dominant, fast rate is higher than the median association rate for yeast eIF4E to a population of yeast mRNAs obtained from cells.^47^ However, it is an order of magnitude slower than for binding of human eIF4E to the m^7^GpppG cap structure analog as measured by rapid-mixing stopped-flow fluorescence spectroscopy (184 µM^−1^ s^−1^).^65^ This is consistent with steric hindrance from the RNA body reducing cap accessibility for eIF4E binding.^30^

The eIF4E•vRNA dissociation rate of 0.93 ± 0.05 s^−1^ (Fig. 2F) was ∼90-fold slower than that measured by stopped-flow for the human eIF4E•m^7^GpppG complex at 100 mM KCl, and ∼11-fold slower than for an eIF4E-complex with an oligoribonucleotide capped with the anti-reverse cap structure analog ([3’-OMe-m^7^G](5’)ppp(5’)G) at 350 mM KCl.^65^ While no structure exists of free eIF4E complexed with a capped RNA oligonucleotide or mRNA, there is evidence for secondary interactions between the RNA body and the dorsal surface of the protein, and these interactions are expected to stabilize the protein-RNA complex by extending its lifetime. Several basic amino acid residues line a groove along this surface,^66,67^ and NMR studies have provided evidence for chemical shift changes in these residues specifically induced by the presence of RNA sequence beyond the cap structure.^68^ The reported stopped-flow kinetics data also led to suggestion of the presence of a second eIF4E–mRNA interaction that impacts the dissociation rate.^65^ Our results are consistent with this proposal.

Taken together, our kinetic data place the eIF4E–UTR equilibrium dissociation constant (*k*_off_ / *k*_on_ = *K*_d_) in the range of ∼74 nM. This represents affinity similar to that determined in past measurements with capped oligonucleotides and mammalian^63^ and yeast^46^ eIF4E.

To establish whether this kinetic signature was particular to the SARS-CoV-2 UTR, or whether it is more general, we also monitored Cy5-eIF4E binding to full-length, polyadenylated human *GAPDH* mRNA. The kinetics for *GAPDH* were similar to the viral UTR, with a double-exponential association-rate distribution (*k*_on_ = 15.8 ± 1.1 µM^−1^ s^−1^ (89%), 2.1 ± 0.5 µM^−1^ s^−1^; *k*_off_ = 0.84 ± 0.1 s^−1^) (Fig. 2E,F). Thus, the eIF4E–UTR kinetic signature appears to resemble eIF4E interaction with a well-translated human mRNA.

Translational activation by phospho-eIF4E^S209^ is at odds with past findings that phospho-eIF4E binds the cap structure and short capped oligonucleotides with lower affinity than non-phosphorylated eIF4E.^62,65^ Thus, it remained unclear whether eIF4E Ser^209^ phosphorylation modulates translation directly through altering the eIF4E–RNA interaction, or instead through an indirect mechanism.

We found that UTR binding of the phosphomimetic variant eIF4E(S209D) was kinetically similar to wild-type eIF4E. The arrival-time distribution remained double-exponential, with association rates of 13.7 ± µM^−1^ s^−1^ and 4.1 ± 0.3 µM^−1^ s^−1^ for the fast and slow phases, respectively (Fig. 2E). The relative amplitude of the fast and slow phases was also similar overall to wild-type eIF4E, with a fast-phase amplitude of ∼71%. The similarity between the slow-phase amplitudes for eIF4E and eIF4E(S209D) points to interaction with different UTR conformations as the source of the double-exponential behaviour. The dissociation rate for eIF4E(S209D) remained unchanged within experimental error relative to wild-type eIF4E, at 0.85 ± 0.11 s^−1^ (Fig. 2F). These rate values lead to an estimated *K*_d_ of ∼62 nM for the eIF4E(S209D)•UTR complex, which is indistinguishable within experimental error from wild-type eIF4E. While the aspartate substitution may be a relatively poor phosphomimetic for binding to the isolated cap structure, it showed weakened interaction with a capped RNA oligonucleotide,^62^ and its overexpression recapitulates the effects of eIF4E phosphorylation in several contexts.^59–61^ We therefore interpret these results to indicate that the mRNA beyond the immediate cap structure plays a role in determining the kinetics of eIF4E–cap interaction, as observed previously for structured oligonucleotides with yeast eIF4E.^46^ This would be consistent with the finding that eIF4E phosphorylation upregulates translation of specific mRNA subsets rather than global translation.^69^

### Human eIF4G(557–1137) does not alter the dynamics of eIF4E–cap interaction

Active eIF4E in cells is found in a complex with the second eIF4F subunit, eIF4G.^67^ Numerous studies have found that eIF4G enhances eIF4E–cap association,^46,70–73^ consistent with a role in promoting translation; the RNA-binding activity of human eIF4G was found to be required for efficient eIF4E–cap binding.^24^ Conversely, other studies have failed to demonstrate an enhancement of eIF4E–cap interaction by eIF4G.^74,75^ Indeed, a stopped-flow kinetic analysis of eIF4E binding to m^7^GpppG showed no change in eIF4E association or dissociation rates with m^7^GpppG in the presence of the eIF4G binding domain for eIF4E.^62^ However, the interaction kinetics were not investigated on full-length mRNAs or mRNA leaders, and the eIF4G fragment used for the stopped-flow studies lacked the RNA-binding region of eIF4G. In parallel, a thermodynamic analysis of human eIF4F–mRNA interaction also indicated that the eIF4G–RNA interaction dominates the free energy of eIF4F–mRNA binding relative to eIF4E–cap binding, implying that the eIF4E•eIF4G dimer has no intrinsic preference for the mRNA 5’ end.^76^ This analysis found a minimal RNA length requirement for robust eIF4F–RNA interaction. Taken together, these results raised the question of whether the eIF4G RNA-binding activity may modulate its ability to influence eIF4E–cap interaction kinetics.

We investigated how a larger eIF4G fragment (residues 557–1137), containing the eIF4E binding domain and the HEAT-repeat region that binds eIF4A, eIF3, and RNA, impacted the kinetics of eIF4E– UTR interaction. This fragment has previously been used in studies of allosteric communication between eIF4E and eIF4A.^28^ We confirmed the presence of the eIF4E- and eIF4A-binding sites in the purified protein by peptide mass mapping (Fig. S2A). The purified eIF4G fragment also stimulated eIF4A ATPase activity in the presence of the SARS-CoV-2 5’ UTR, to a similar or even greater degree than reported previously for other RNAs (Fig. S2B).^28^

eIF4G(557–1137) bound the viral UTR in a highly cooperative manner (Fig. 3A), as assessed by electrophoretic mobility shift assay, and as observed previously for the full eIF4F complex.^76^Stoichiometric binding was saturated at 75 nM eIF4G fragment, while additional eIF4G molecules were bound at 300 nM, resulting in a clear supershift of the RNA band. This indicates high-affinity RNA-protein interaction, consistent with the previously reported *K*_½,app_ value of ∼17 nM for eIF4F–mRNA interaction in the absence of ATP.^76^ This affinity also implies that the viral UTR will be substantially bound at a 1:1 stoichiometry by the eIF4G fragment under the conditions of the single-molecule experiments described below.

**Figure 3.**
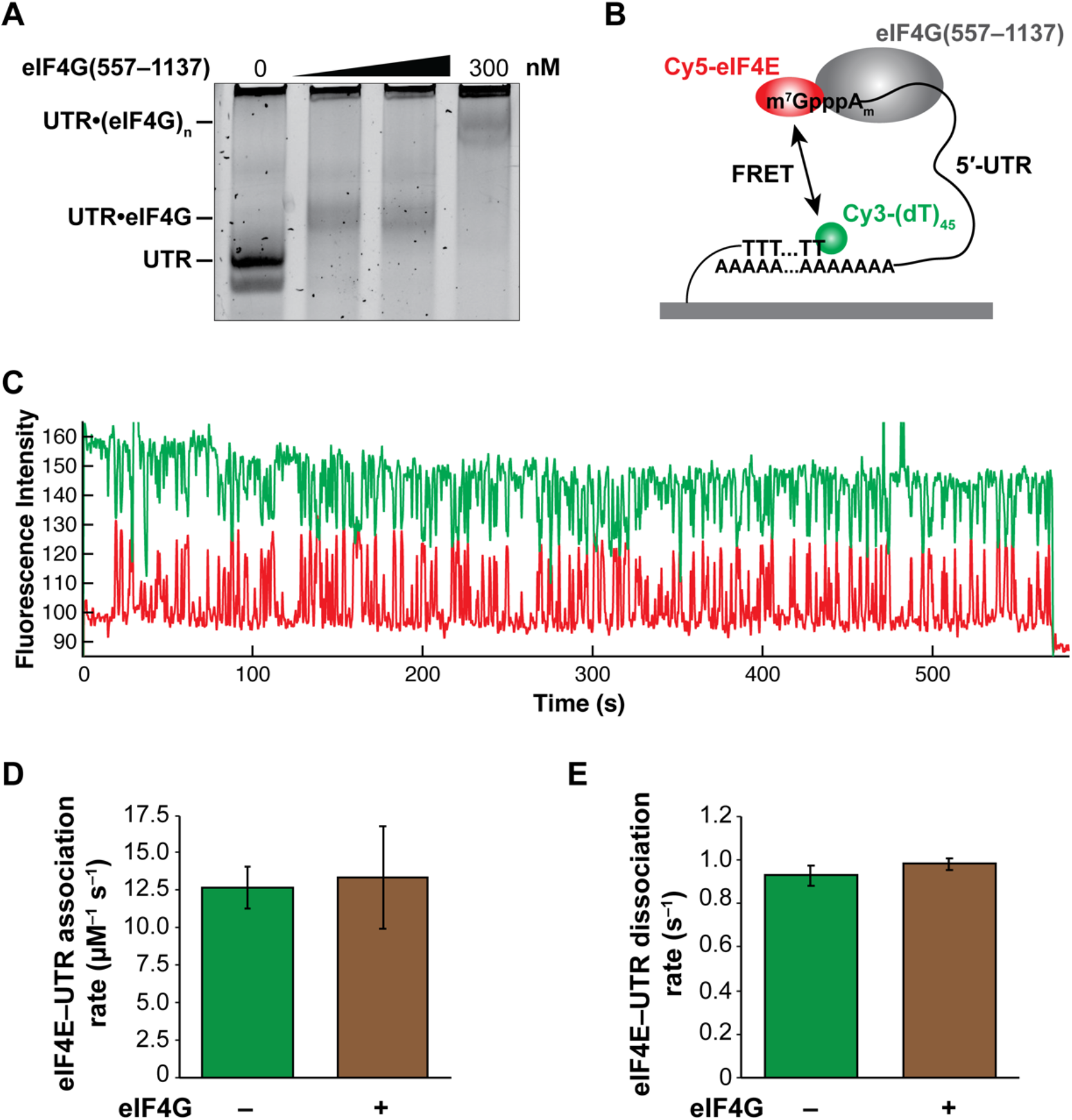
Effects of eIF4G(557–1137) on eIF4E interaction with the viral UTR. (A) Electrophoretic mobility shift assay for eIF4G–UTR binding, carried out by nondenaturing PAGE. The UTR concentration was ∼25 nM. (B) Schematic of single-molecule fluorescence experiment including the eIF4G fragment. (C) Representative single-molecule fluorescence trace for eIF4E–UTR interaction in the presence of eIF4G(557–1137). (D) eIF4E– and eIF4E(S209D)–UTR association rates in the presence of eIF4G(557–1137). (E) eIF4E– and eIF4E(S209D)–UTR dissociation rates in the presence of eIF4G(557–1137).

We then co-delivered Cy5-eIF4E (30 nM) to the surface-immobilized UTR in the presence of eIF4G(557–1137) (40 nM; Fig. 3B). We again observed robust eIF4E–UTR FRET (Fig. 3C). The eIF4E– RNA association rate was 13.4 ± 3.5 µM^−1^ s^−1^ (Fig. 3D), which was not significantly different to the rate observed with eIF4E alone. However, the slow-phase binding interaction was absent in the presence of the eIF4G fragment, i.e., the arrival-time distribution became essentially single-exponential (Supplementary Table 1). This is consistent with the eIF4G RNA-binding activity modulating the UTR conformational ensemble to bias it toward a configuration that is more accessible for eIF4E binding.

Meanwhile, the eIF4E–UTR dissociation rate also remained essentially unchanged in the presence of eIF4G(557–1137), at 0.98 ± 0.03 s^−1^ (Fig. 3E). Thus, while we cannot rule out that the situation for full-length eIF4G may be different, our results indicate that eIF4G–RNA binding does not substantially alter the kinetics of eIF4E–cap interaction for a given RNA conformation. This mirrors the findings from the previous stopped-flow studies.

### Free eIF4A accelerates the eIF4E–UTR association rate

eIF4A is an ATP-dependent RNA helicase, and the archetypal member of the DEAD-box RNA helicase family.^13^ Incorporation into the eIF4F complex activates the eIF4A helicase and ATPase activities, including by allosteric communication between eIF4E and eIF4A through eIF4G.^28^ However, because eIF4A is expressed at much higher levels than eIF4E or eIF4G in cells,^77^ the majority of the enzyme is not eIF4F-bound. A systems-biology analysis in yeast suggested that the free eIF4A fraction also plays a significant role in promoting translation, since small reductions in eIF4A abundance – to an extent not expected to deplete active eIF4F – resulted in a linear decrease in translational output, in proportion to the degree of eIF4A depletion.^78^

The SARS-CoV-2 5’-UTR contains three conserved stem loop structures.^79^ In part these stem loops promote translation through conferring resistance to endonucleolytic cleavage by the viral nsp1 protein, which cleaves host mRNAs during its interaction with the mRNA entry channel of the 40S ribosomal subunit.^79–81^ While the exact function of these structures in viral translation is not well understood, it seems more than likely that helicase activity would be required to resolve them during viral translation initiation. Interestingly, the start codon is embedded in one of the stem loops.^38^

We recently showed that free yeast eIF4A accelerated binding of yeast eIF4E to mRNA, i.e., independently of its incorporation in the eIF4F complex.^47^ However, yeast and human eIF4E have significantly different affinities for single- and double-stranded RNA, and different preferences for RNA overhangs needed to load the helicase onto the RNA.^82,83^ Thus, it remained unclear whether free human eIF4A would have a similar effect, and whether any effect would be significant for the SARS-CoV-2 RNA.

Addition of 1 µM eIF4A with 1 mM ATP doubled the eIF4E–UTR binding rate, to 25.4 ± 0.7 µM^−1^ s^−1^ (Fig. 4A,B). The arrival-time distribution again became essentially single-exponential (slow-phase amplitude ∼1%; Supplementary Table 1). The eIF4E–UTR dissociation rate remained unchanged relative to the eIF4E-only condition, at 0.87 ± 0.08 s^−1^ (Fig. 4C). Acceleration was also observed for eIF4E(S209D); however, the effect was less pronounced at only a ∼37% increase in *k*_on_, to 18.6 ± 1.1 µM^−1^ s^−1^ (Fig. 4B), with a single-exponential arrival-time distribution. eIF4A also did not alter the eIF4E(S209D) dissociation rate, which remained at 0.80 ± 0.04 s^−1^ (Fig. 4C).

**Figure 4.**
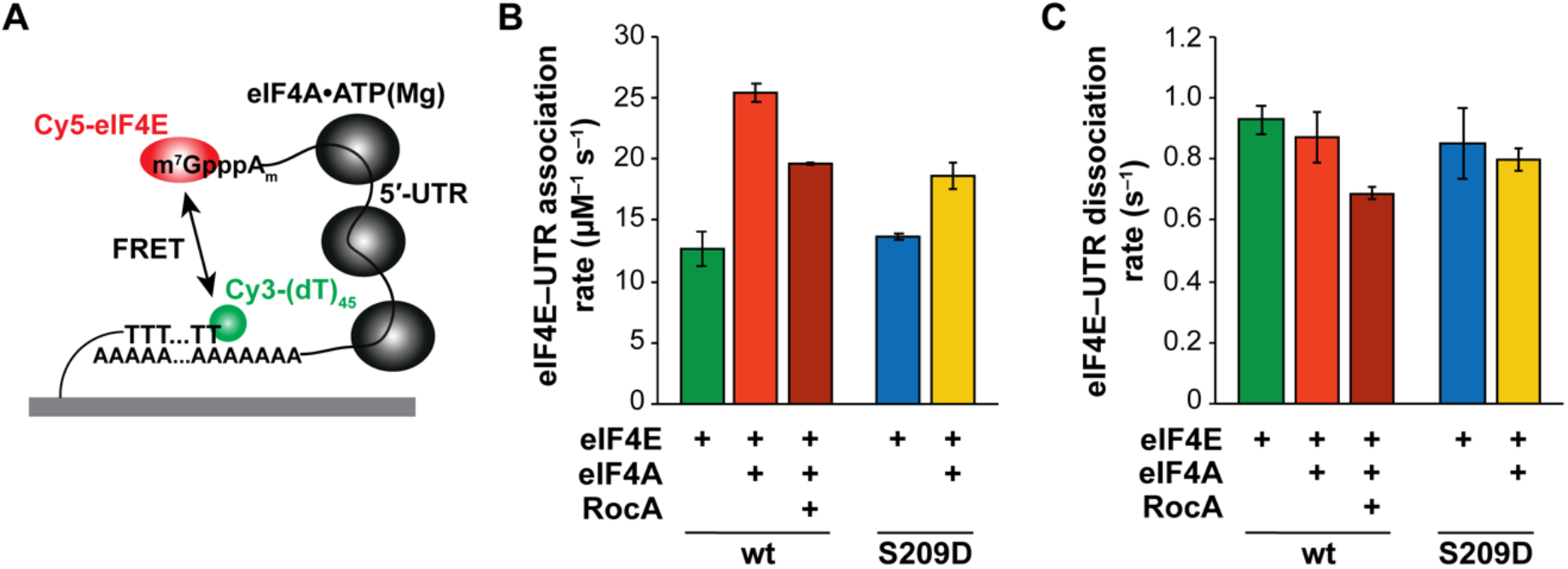
Effects of free eIF4A•ATP(Mg) on eIF4E–UTR interaction. (A) Schematic of single-molecule fluorescence experiment including free eIF4A. (B) eIF4E– and eIF4E(S209D)–UTR association rates in the presence of 1 µM eIF4A with 1 mM ATP•Mg, along with the effect of RocA (20 µM) on eIF4E(wt)– UTR association rate. (C) eIF4E– and eIF4E(S209D)–UTR dissociation rates in the presence of free eIF4A.

Thus, the effect of eIF4A was to increase eIF4E–cap affinity, entirely through effects on association. While eIF4A requires ATP to bind tightly to RNA,^84^ the enzyme is an ineffective helicase without activation from its cofactors, i.e. eIF4G and eIF4B/eIF4H (Fig. S2B and Ref. 85). In accord with our data for yeast eIF4A,^47^ we interpret the increase in eIF4E–UTR association rate, without a change to the dissociation rate, as resulting from eIF4A reducing steric block to cap accessibility. Since significant eIF4A helicase activity is unlikely under our experimental conditions without eIF4G present (Fig. S2B), the simplest model is that eIF4A–RNA binding disrupts tertiary contacts in the UTR, then clamps the UTR in a conformation where the cap structure is more accessible to eIF4E. This would be consistent with the RNA-clamping activity that has been characterized for other DEAD-box helicases.^86^ These results suggest a model where, once RNA tertiary compaction is released by eIF4A–UTR binding, the eIF4E binding rate is then controlled by secondary structures near the 5’ end,^46,47^ but the effect is dampened by the negative charge at eIF4E^Asp209^, resulting in the slower degree of acceleration observed for eIF4E(S209D).

The rocaglate eIF4A inhibitor RocA potently reduced translation of our luciferase reporter RNA (Fig. 1C), in agreement with past findings on rocaglate inhibition of HCoV-229E and MERS-CoV translation that attenuates viral replication,^39,40^ and recent findings on inhibition of SARS-CoV-2 translation by the synthetic rocaglate CR-31-B(–).^56^ However, two modes of action have been proposed for translation inhibition by rocaglates – blockage of scanning due to locking of eIF4A onto mRNA leaders,^52^ and prolonged eIF4F–mRNA interaction, induced by inhibition of eIF4F-bound eIF4A,^54^ which exerts a *trans*-negative effect on translation initiation on other mRNAs.

Given these potential modes of action, we asked whether RocA impacts the ability of free eIF4A to modulate eIF4E–UTR interaction dynamics. We monitored eIF4E–UTR binding in the presence of eIF4A, ATP, and 20 µM RocA, approximately 40 times the RocA concentration which reduced reporter translation to near-baseline levels, and within the concentration range where RocA induced eIF4F–cap sequestration.^54^ The eIF4E–UTR binding rate was reduced by RocA addition, at 19.6 ± 0.1 µM^−1^ s^−1^ (Fig. 4B) and the arrival-time distribution remained essentially single-exponential. However, the dissociation rate decreased slightly, to 0.69 ± 0.02 s^−1^ (Fig. 4C). This was the sole condition where we observed significantly prolonged eIF4E–UTR interaction in this study. We note that the binding and dissociation rates were decreased to the same extent, consistent with clamping of eIF4A onto the UTR directly modulating eIF4E–UTR interaction.

### Coordinated activities of eIF4G and eIF4A determine the eIF4E–cap binding rate in the eIF4F complex

We next characterized how formation of the entire eIF4F complex impacted eIF4E–UTR interaction dynamics. To the surface-immobilized viral UTR we thus co-delivered Cy5-eIF4E (30 nM), along with eIF4G(557–1137) (40 nM) and eIF4A (1 µM), to recapitulate physiologically-relevant relative factor concentrations. Experiments were carried out in the presence of 1 mM Mg•ATP (Fig. 5A).

**Figure 5.**
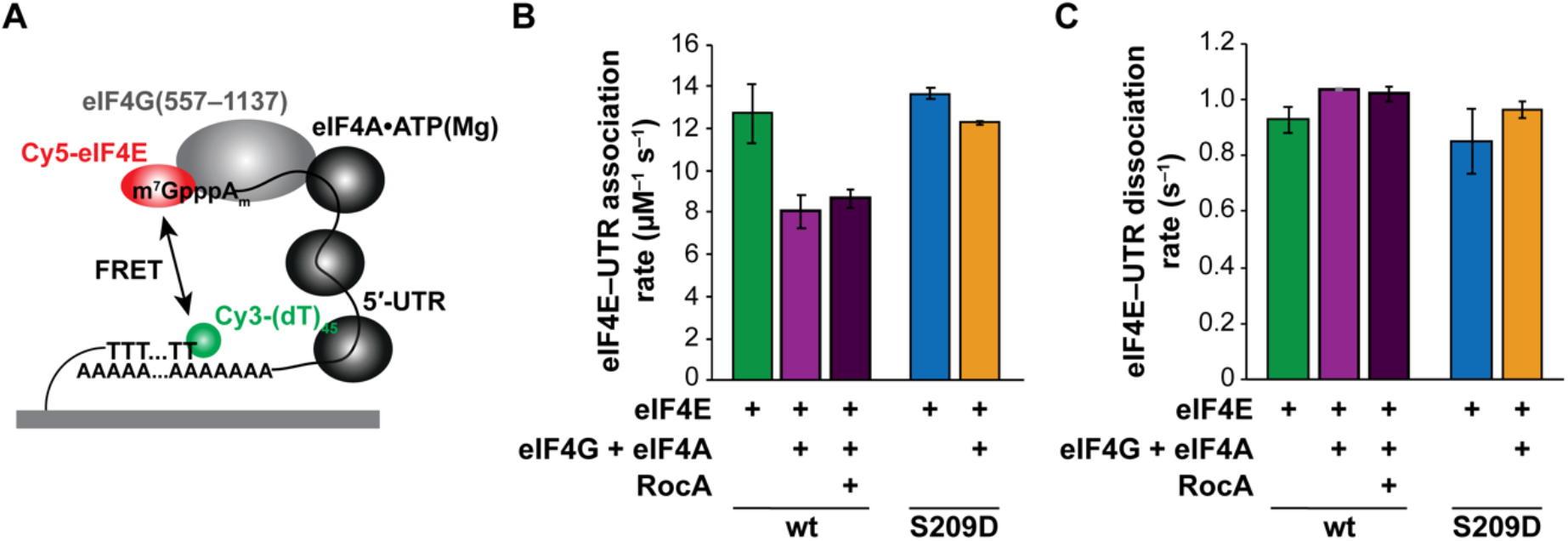
eIF4E–UTR interaction in the eIF4F complex. (A) Schematic of single-molecule fluorescence experiment including eIF4G(557–1137) (40 nM) and eIF4A (1 µM) along with Cy5-eIF4E (30 nM). (B) eIF4E– and eIF4E(S209D)–UTR association rates in the presence of eIF4G(557–1137) and eIF4A with 1 mM ATP•Mg, along with the effect of RocA (20 µM) on kinetics of the eIF4F complex reconstituted with wild-type eIF4E. (C) eIF4E– and eIF4E(S209D)–UTR dissociation rates in the presence of eIF4G(557–1137), eIF4A and ATP.

Surprisingly, eIF4E–UTR binding slowed substantially in the presence of the eIF4F complex relative to all other conditions. This entirely reversed the on-rate enhancement by free eIF4A, even though free eIF4A in this experiment was still present in order-of-magnitude excess over the eIF4F complex. The association rate was 8.1 ± 0.8 µM^−1^ s^−1^, a reduction of 1.6-fold relative to eIF4E alone, and 3.1-fold relative to eIF4E binding in the presence of free eIF4A alone (Fig. 5B). The arrival-time distribution was again single-exponential (Supplementary Table 1). The eIF4E dissociation rate was again essentially unchanged, at 1.03 ± 0.03 s^−1^ (Fig. 5C). Addition of RocA did not significantly alter these kinetics, with association and dissociation rates of 8.66 ± 0.47 µM^−1^ s^−1^ and 1.02 ± 0.03 s^−1^, respectively (Fig. 5B,C). Combining our luciferase reporter results with these single-molecule kinetics, we conclude that RocA inhibits translation driven by the SARS-CoV-2 UTRs at a point after cap recognition, most likely during ribosomal scanning.^52^

These rates suggest a *K*_d_ value of ∼127 nM for the eIF4F•UTR complex. This represents seven-fold lower affinity than binding of native eIF4F to the capped β-globin mRNA measured by electrophoretic mobility shift in the presence of ATP,^76^ a result that is consistent with impairment of eIF4F–mRNA binding by secondary structure in the 5’ UTR.

Reduced frequency of eIF4E–UTR binding events raised the possibility that the helicase activity of eIF4A, which is activated by eIF4G, caused RNA structural remodelling that increased the RNA end-to-end distance beyond the FRET range. However, two lines of evidence argue against this possibility. First, we did not observe molecules that showed frequent FRET at the beginning of the movie, but where FRET disappeared during movie acquisition. This behavior would be expected if eIF4A disrupted the RNA:DNA hybrid duplex immobilizing the UTR. Second, imaging of Cy5-eIF4E by direct excitation after prolonged incubation with eIF4F and ATP did not reveal the presence of eIF4E–UTR binding events occurring without FRET (Fig. S3). Thus, the reduced eIF4E–UTR binding rate with eIF4F and ATP appears to be a *bona fide* outcome of forming the full eIF4F complex. We also note here that this result provides further validation that our FRET signal yields a good measure of the rate constants for eIF4E binding and dissociation. Lack of eIF4E binding in the eIF4F complex without FRET also points to an intrinsic and strong 5’-end bias for eIF4E–mRNA binding in the eIF4F complex.

Loss of the acceleration of eIF4E–cap association afforded by free eIF4A when eIF4E was bound in the eIF4F heterotrimer suggested that steric block to physical eIF4E interaction with the cap structure no longer limited the binding rate. One explanation for this result is that cap binding of the full eIF4F complex is limited by a different process than binding of free eIF4E or eIF4E•eIF4G. Such a model agrees overall with the proposal that eIF4E plays a minimal role in determining eIF4F–mRNA affinity.^76^ In the previous study the division of labor between eIF4G- and eIF4A-RNA binding in stabilizing eIF4F– mRNA interactions was not delineated, though eIF4G was favored as the eIF4F subunit that dominated the binding affinity. Our results suggest that both eIF4G and eIF4A contribute, and that direct eIF4G•eIF4A interaction is required to establish the net eIF4F–mRNA affinity. In support of this model, we note that for two of three mRNAs studied in the previous work,^76^ addition of ATP reduced eIF4F•mRNA affinity by a similar extent to the reduction we observed in eIF4E–cap binding rates with eIF4F•ATP relative to eIF4E•eIF4G alone. An alternative but not mutually exclusive model is that free eIF4A bound distributively throughout the UTR acts as a competitive inhibitor of distributive eIF4F binding, except at the mRNA cap where an additional point of eIF4F–UTR attachment is available, i.e., the cap structure. This would rationalize the observed apparent 5’-end bias for eIF4F–UTR binding.

However, the situation for the eIF4F complex formed with eIF4E(S209D) was significantly different. With this variant, only a slight deceleration in eIF4E–UTR association was observed relative to free eIF4E(S209D) with or without eIF4G(557–1137), though the rate was again reduced relative to eIF4E(S209D) with free eIF4A. The association rate was 12.3 ± 0.1 µM^−1^ s^−1^, and the dissociation rate was 0.96 ± 0.03 s^−1^ (Fig. 5B,C). The net outcome of this effect was that the phosphomimetic eIF4F complex had a greater apparent affinity for the capped UTR, with an estimated *K*_d_ ∼ 78 nM. This finding may offer an explanation for why binding to the cap structure or to capped RNAs of eIF4E(S209D), and by implication phospho-eIF4E, is weaker than non-phosphorylated eIF4E, whilst eIF4E phosphorylation promotes translation initiation – since translation initiation is mediated by the intact eIF4F complex. In this model, eIF4E phosphorylation modulates or abrogates the allosteric crosstalk between the eIF4F subunits, resulting in a greater net eIF4F–UTR affinity.

In summary, our results provide a real-time characterization of eIF4E interaction dynamics with a full-length mRNA 5’-untranslated region, along with a kinetic dissection of the contributions of each eIF4F subunit to cap binding. The kinetic data suggest a role for free eIF4A in significantly remodelling mRNA tertiary structure to modulate cap accessibility. Such interactions are consistent with distributive mRNA binding of free eIF4A playing an important function in early translation initiation,^87^ and consistent with acceleration of pre-initiation complex recruitment by yeast eIF4A due to eIF4A–mRNA interactions remote from the cap structure.^88^ In turn, the results support a mechanism where rocaglate eIF4A inhibitors may potently suppress viral translation through blocking ribosomal scanning. Our data indicate that coordination of eIF4G and eIF4A activities in the eIF4F complex play a deterministic role in establishing the efficiency of cap recognition. They also suggest that the phosphorylation status of eIF4E can be sensed by the two other eIF4F subunits, echoing the ability of eIF4E to allosterically communicate with eIF4A through eIF4G.^28^ Taken together, these results highlight that a complex interplay of eIF4F subunit activities and mRNA structure, likely tunable by post-translational modification of eIF4E, plays a central role in determining the efficiency of early translation initiation. Our findings also point to potential benefits of rationally designing 5’ UTR sequences in mRNA-based vaccines and therapeutics.

## AVAILABILITY

All data generated and analysed in the present study are included in the manuscript and supplementary information.

## SUPPLEMENTARY DATA

Supplementary Data are available at NAR online.

## ACKNOWLEDGEMENTS

The pULTRA plasmid and *E. coli* BL95ΔAΔ*fabR* strain were generous gifts of Abhishek Chatterjee (Boston College). We thank Jin Chen (UT Southwestern Medical Center) and Gregor Blaha (UC Riverside) for insightful discussions during preparation of the manuscript.

