## Supplementary Information for "Single-Molecule Dynamics of SARS-CoV-2 5’ Cap Recognition by Human eIF4F"

Hea Jin Hong[1], Matthew G. Guevara[1], Eric Lin[1], and Seán E. O'Leary[1]
[1] Department of Biochemistry, University of California Riverside, Riverside, CA 92521, U.S.A.

### TABLE AND FIGURE LEGENDS

Supplementary Table 1. Rate constants, fast- and slow-phase association-rate amplitudes, and numbers of molecules and events analyzed for each experimental condition

Figure S1. Reagent preparation and validation. (A) SDS-PAGE analysis of recombinant proteins. (B) Size-exclusion chromatogram of purified Cy5-eIF4E, with traces for protein absorbance (280 nm) and Cy5 absorbance (645 nm). (C) Size-exclusion chromatogram of purified Cy5-eIF4E(S209D).

Figure S2. Validation of eIF4G(557–1137) activity. (A) Peptide mass-mapping data indicating presence of the eIF4E and eIF4A-binding domains. (B) Stimulation by eIF4G(557–1137) of eIF4A ATPase activity with the SARS-CoV-2 5' UTR.

Figure S3. Direct detection of Cy5-eIF4E–UTR binding events by 642-nm Cy5 excitation does not reveal binding events occurring without FRET in the presence of eIF4A and eIF4G. (A) Schematic of dual-illumination experiment. (B) Representative single-molecule fluorescence trace from dual illumination of Cy3-UTR and Cy5-eIF4E (30 nM) in the presence of unlabelled eIF4G (40 nM) and eIF4A (1 μM), with 1 mM ATP.

### Supplementary Table 1

| Experiment – SARS-CoV-2 UTR | k_on,1 1/(µM s) Fast arrival | Fast-phase amplitude (%) | k_on,2 1/(µM s) Slow arrival | Slow-phase amplitude (%) | k_off (1/s) | No. of molecules | No. of events |
|---|---|---|---|---|---|---|---|
| hs4E-WT (30 nM) (1) | 13.69 | 77.10 | 2.95 | 22.90 | 0.90 | 165 | 9296 |
| hs4E-WT (30 nM) (2) | 11.73 | 86.10 | 3.21 | 13.90 | 0.96 | 99 | 7119 |
| Average | 12.71 | 81.60 | 3.08 | 18.40 | 0.93 | | |
| stdev | 1.39 | 6.36 | 0.18 | 6.36 | 0.05 | | |
| hs4E-WT (30 nM) + hs4A (1 uM) (1) | 25.84 | 98.89 | 1.50 | 1.11 | 0.93 | 167 | 24104 |
| hs4E-WT (30 nM) + hs4A (1 uM) (2) | 24.85 | 98.06 | 1.84 | 1.94 | 0.81 | 137 | 18259 |
| Average | 25.35 | 98.48 | 1.67 | 1.53 | 0.87 | | |
| stdev | 0.70 | 0.59 | 0.24 | 0.59 | 0.08 | | |
| hs4E-WT (30 nM) + hs4G (40 nM) (1) | 15.81 | 97.76 | 1.752 | 2.24 | 1.00 | 136 | 13712 |
| hs4E-WT (30 nM) + hs4G (40 nM) (2) | 10.93 | 99.15 | 0.712 | 0.85 | 0.96 | 129 | 9659 |
| Average | 13.37 | 98.45 | 1.23 | 1.55 | 0.98 | | |
| stdev | 3.45 | 0.98 | 0.74 | 0.98 | 0.03 | | |
| hs4E-WT (30 nM) + hs4G (40 nM) + hs4A (1 uM) (1) | 7.513 | 98.84 | 0.48 | 1.16 | 1.04 | 77 | 3464 |
| hs4E-WT (30 nM) + hs4G (40 nM) + hs4A (1 uM) (2) | 8.613 | 99.00 | 0.52 | 1.00 | 1.03 | 80 | 5048 |
| Average | 8.06 | 98.92 | 0.50 | 1.08 | 1.03 | | |
| stdev | 0.78 | 0.12 | 0.03 | 0.12 | 0.00 | | |
| hs4E-WT (30 nM) + hs4G (40 nM) + hs4A (1 uM) + RocA (20 uM) (1) | 8.32 | 90.00 | 2.45 | 10.00 | 1.00 | 76 | 3406 |
| hs4E-WT (30 nM) + hs4G (40 nM) + hs4A (1 uM) + RocA (20 uM) (2) | 8.99 | 96.20 | 1.22 | 3.80 | 1.04 | 93 | 3558 |
| Average | 8.66 | 93.10 | 1.83 | 6.90 | 1.02 | | |
| stdev | 0.47 | 4.38 | 0.87 | 4.38 | 0.03 | | |
| hs4E-WT (30 nM) + hs4A (1 uM) + RocA (20 uM) (1) | 19.53 | 97.24 | 2.03 | 2.76 | 0.67 | 105 | 11010 |
| hs4E-WT (30 nM) + hs4A (1 uM) + RocA (20 uM) (2) | 19.66 | 99.41 | 0.52 | 0.59 | 0.70 | 127 | 10459 |
| Average | 19.60 | 98.33 | 1.28 | 1.68 | 0.69 | | |
| stdev | 0.09 | 1.53 | 1.07 | 1.53 | 0.02 | | |
| hs4E-S209D (30 nM) (1) | 13.82 | 66.6 | 3.84 | 33.4 | 0.9302 | 86 | 2872 |
| hs4E-S209D (30 nM) (2) | 13.48 | 76.2 | 4.25 | 23.8 | 0.7695 | 71 | 1244 |
| Average | 13.65 | 71.40 | 4.05 | 28.60 | 0.85 | | |
| stdev | 0.25 | 6.79 | 0.29 | 6.79 | 0.11 | | |
| hs4E-S209D (30 nM) + hs4A (1 uM) (1) | 17.90 | 97.6 | 1.795 | 2.4 | 0.8228 | 107 | 7977 |
| hs4E-S209D (30 nM) + hs4A (1 uM) (2) | 18.13 | 98.9 | 0.679 | 1.1 | 0.7586 | 43 | ND |
| hs4E-S209D (30 nM) + hs4A (1 uM) (3) | 19.87 | 98.86 | 1.185 | 1.14 | 0.8188 | 101 | 2615 |
| Average | 18.64 | 98.45 | 1.22 | 1.55 | 0.80 | | |
| stdev | 1.08 | 0.74 | 0.56 | 0.74 | 0.04 | | |
| hs4E-S209D (30 nM) + hs4G (40 nM) + hs4A (1 uM) (1) | 12.19 | 96.1 | 2.13 | 3.9 | 0.9834 | 94 | 6230 |
| hs4E-S209D (30 nM) + hs4G (40 nM) + hs4A (1 uM) (2) | 12.327 | 98.9 | 0.655 | 1.1 | 0.9394 | 92 | 6640 |
| Average | 12.26 | 97.52 | 1.39 | 2.48 | 0.96 | | |
| stdev | 0.09 | 2.00 | 1.05 | 2.00 | 0.03 | | |

| Experiment – GAPDH | k_on,1 1/(µM s) Fast arrival | Fast-phase amplitude (%) | k_on,2 1/(µM s) Slow arrival | Slow-phase amplitude (%) | k_off (1/s) | No. of molecules | No. of events |
|---|---|---|---|---|---|---|---|
| hs4E-WT (30 nM) - GAPDH (1) | 16.55 | 87.7 | 2.49 | 12.3 | 0.77 | 75 | 5015 |
| hs4E-WT (30 nM) - GAPDH (2) | 14.95 | 91.7 | 1.75 | 8.3 | 0.91 | 94 | 5906 |
| Average | 15.75 | 89.70 | 2.12 | 10.30 | 0.84 | | |
| stdev | 1.14 | 2.83 | 0.52 | 2.83 | 0.10 | | |

.

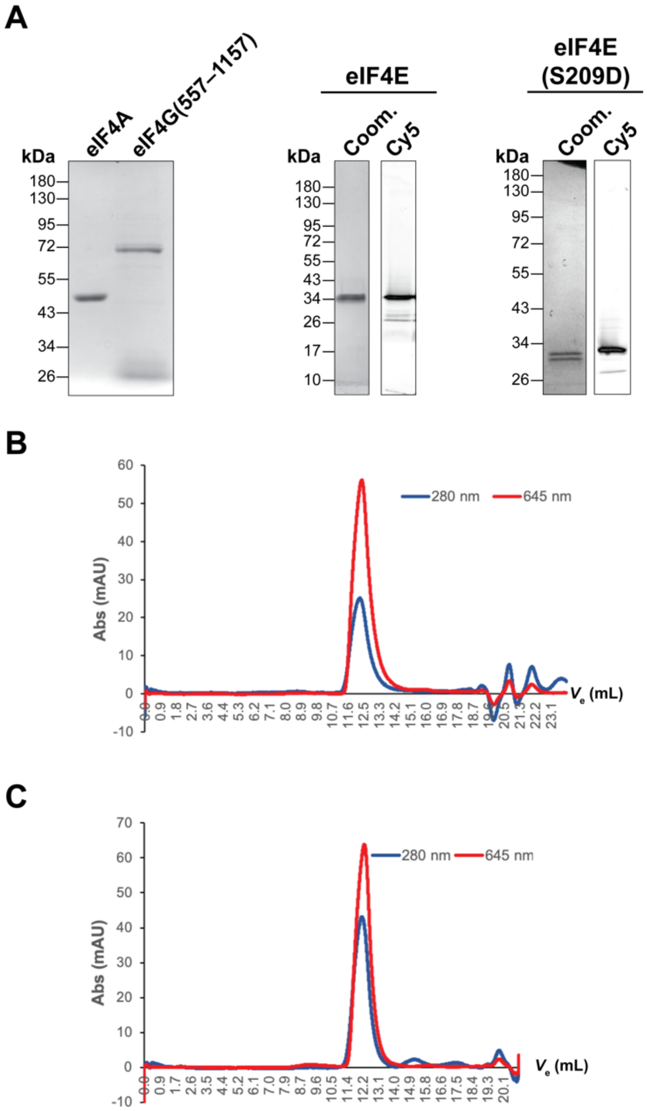
A
eIF4A
eIF4G(557–1157)
kDa
180
130
95
72
55
43
34
26
eIF4E
Coom.
Cy5
17
10
eIF4E (S209D)
B
280 nm
645 nm
Abs (mAU)
$V_e$ (mL)
C

**Figure S1.**

A

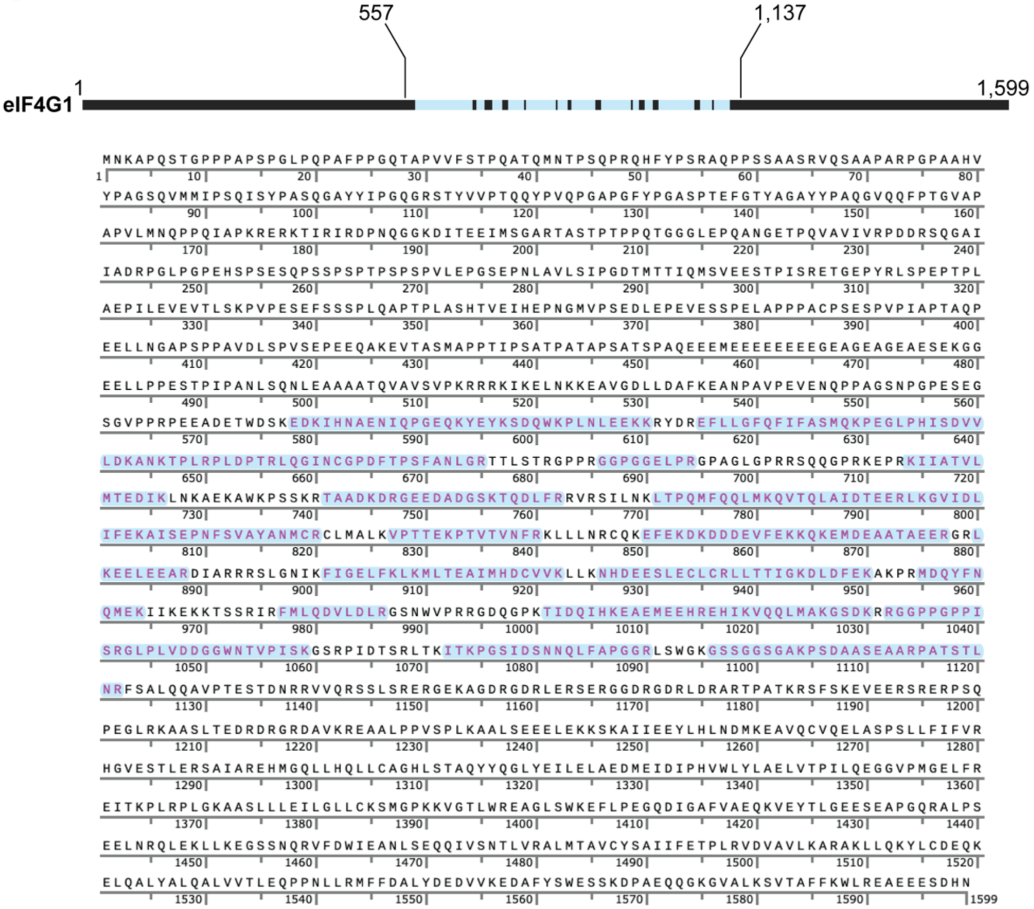
557
1,137
eIF4G1
1
1,599
MNKAPQSTGPPPAPSPGLPQPAFPPGQTAPVVFSTPQATQMNTPSQPRQHFYPSRAQPPSSAASRVQSAAPARPGPAAHV
YPAGSQVMMIPSQISYPASQGAYYIPGQGRSTYVVPTQQYPVQPGAPGFYPGASPTEFGTYAGAYYPAQGVQQFPTGVAP
APVLMNQPPQIAPKRERKTIRIRDPNQGGKDITEEIMSGARTASTPTPPQTGGGLEPQANGETPQVAVIVRPDDRSQGAI
IADRPGLPGPEHSPSESQPSSPSPTPSPSPVLEPGSEPNLAVLSIPGDTMTTIQMSVEESTPISRETGEPYRLSPEPTPL
AEPILEVEVTLSKPVPESEFSSSPLQAPTPLASHTVEIHEPNGMVPSEDLEPEVESSPELAPPPACPSESPVPIAPTAQP
EELLNGAPSPPAVDLSPVSEPEEQAKEVTASMAPPTIPSATPATAPSATSPAQEEEMEEEEEEEEGEAGEAGEAESEKGG
EELLPPESTPIPANLSQNLEAAAATQVAVSVPKRRRKIKELNKKEAVGDLLDAFKEANPAVPEVENQPPAGSNPGPESEG
SGVPPRPEEADETWDSKEDKIHNAENIQPGEQKYEYKSDQWKPLNLEEKKRYDREFLLGFQFIFASMQKPEGLPHISDVV
LDKANKTPLRPLDPTRLQGINCGPDFTPSFANLGRTTLSTRGPPRGGPGGELPRGPAGLGPRRSQQGPRKEPRKIIATVL
MTEDIKLNKAEKAWKPSSKRTAADKDRGEEDADGSKTQDLFRRVRSILNKLTPQMFQQLMKQVTQLAIDTEERLKGVIDL
IFEKAISEPNFSVAYANMCRCLMALKVPTTEKPTVTVNFRKLLLNRCQKEFEKDKDDDEVFEKKQKEMDEAATAEERGRL
KEELEEARDIARRRSLGNIKFIGELFKLKMLTEAIMHDCVVKLLKNHDEESLECLCRLLTTIGKDLDFEKAKPRMDQYFN
QMEKIIKEKKTSSRIRFMLQDVLDLRGSNWVPRRGDQGPKTIDQIHKEAEMEEHREHIKVQQLMAKGSDKRRGGPPGPPI
SRGLPLVDDGGWNTVPISKGSRPIDTSRLTKITKPGSIDSNNQLFAPGGRLSWGKGSSGGSGAKPSDAASEAARPATSTL
NRFSALQQAVPTESTDNRRVVQRSSLSRERGEKAGDRGDRLERSERGGDRGDRLDRARTPATKRSFSKEVEERSRERPSQ
PEGLRKAASLTEDRDRGRDAVKREAALPPVSPLKAALSEEELEKKSKAIIEEYLHLNDMKEAVQCVQELASPSLLFIFVR
HGVESTLERSAIAREHMGQLLHQLLCAGHLSTAQYYQGLYEILELAEDMEIDIPHVWLYLAELVTPILQEGGVPMGELFR
EITKPLRPLGKAASLLLEILGLLCKSMGPKKVGTLWREAGLSWKEFLPEGQDIGAFVAEQKVEYTLGEESEAPGQRALPS
EELNRQLEKLLKEGSSNQRVFDWIEANLSEQQIVSNTLVRALMTAVCYSAIIFETPLRVDVAVLKARAKLLQKYLCDEQK
ELQALYALQALVVTLEQPPNLLRMFFDALYDEDVVKEDAFYSWESSKDPAEQQGKGVALKSVTAFFKWLREAEEESDHN

B

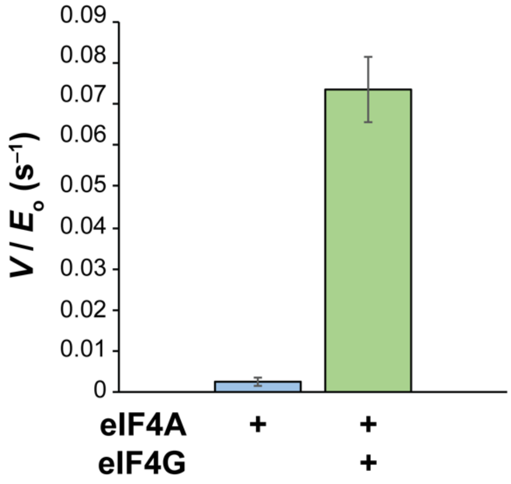
0.09
0.08
0.07
0.06
0.05
0.04
0.03
0.02
0.01
0
V / E$_o$ (s$^{-1}$)
eIF4A
eIF4G
+
+
+

**Figure S2**

**A**

**B**

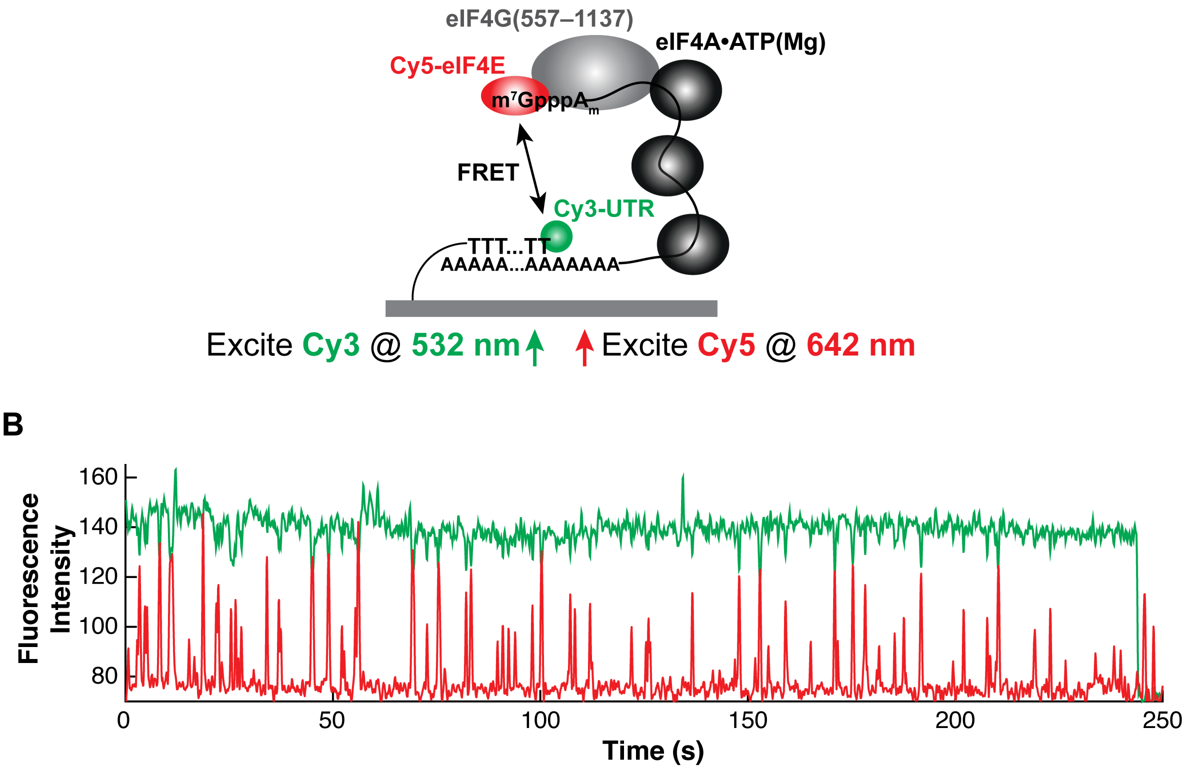

eIF4G(557–1137)
eIF4A•ATP(Mg)
Cy5-eIF4E
m7GpppAm
FRET
Cy3-UTR
TTT...TT
AAAAA...AAAAAAA
Excite Cy3 @ 532 nm
Excite Cy5 @ 642 nm
Fluorescence Intensity
160
140
120
100
80
0
50
100
150
200
250
Time (s)


**Figure S3**

**Insert DNA sequences for RNA template and protein overexpression constructs.**

**SalI-T7-*GAPDH*-EcoRI** (From *Homo sapiens*, transcript variant 1; NM_002406.7)

GTCGACTAATACGACTCACTATAGGTGTTCGACAGTCAGCCGCATCTTCTTTTGCGTCGCCAGC
CGAGCCACATCGCTCAGACACCATGGGGAAGGTGAAGGTCGGAGTCAACGGATTTGGTCGTATT
GGGCGCCTGGTCACCAGGGCTGCTTTTAACTCTGGTAAAGTGGATATTGTTGCCATCAATGACC
CCTTCATTGACCTCAACTACATGGTTTACATGTTCCAATATGATTCCACCCATGGCAAATTCCA
TGGCACCGTCAAGGCTGAGAACGGGAAGCTTGTCATCAATGGAAATCCCATCACCATCTTCCAG
GAGTGAGTGGAAGACAGAATGGAAGAAATGCGAGATCCCTCCAAAATCAAGTGGGGCGATGCTG
GCGCTGAGTACGTCGTGGAGTCCACTGGCGTCTTCACCACCATGGAGAAGGCTGGGGCTCATTT
GCAGGGGGGAGCCAAAAGGGTCATCATCTCTGCCCCCTCTGCTGATGCCCCCATGTTCGTCATG
GGTGTGAACCATGAGAAGTATGACAACAGCCTCAAGATCATCAGCAATGCCTCCTGCACCACCA
ACTGCTTAGCACCCCTGGCCAAGGTCATCCATGACAACTTTGGTATCGTGGAAGGACTCATGAC
CACAGTCCATGCCATCACTGCCACCCAGAAGACTGTGGATGGCCCCTCCGGGAAACTGTGGCGT
GATGGCCGCGGGGCTCTCCAGAACATCATCCCTGCCTCTACTGGCGCTGCCAAGGCTGTGGGCA
AGGTCATCCCTGAGCTGAACGGGAAGCTCACTGGCATGGCCTTCCGTGTCCCCACTGCCAACGT
GTCAGTGGTGGACCTGACCTGCCGTCTAGAAAAACCTGCCAAATATGATGACATCAAGAAGGTG
GTGAAGCAGGCGTCGGAGGGCCCCCTCAAGGGCATCCTGGGCTACACTGAGCACCAGGTGGTCT
CCTCTGACTTCAACAGCGACACCCACTCCTCCACCTTTGACGCTGGGGCTGGCATTGCCCTCAA
CGACCACTTTGTCAAGCTCATTTCCTGGTATGACAACGAATTTGGCTACAGCAACAGGGTGGTG
GACCTCATGGCCCACATGGCCTCCAAGGAGTAAGACCCCTGGACCACCAGCCCCAGCAAGAGCA
CAAGAGGAAGAGAGAGACCCTCACTGCTGGGGAGTCCCTGCCACACTCAGTCCCCCACCACACT
GAATCTCCCCTCCTCACAGTTGCCATGTAGACCCCTTGAAGAGGGGAGGGGCCTAGGGAGCCGC
ACCTTGTCATGGAATTC

**AscI-T7-SARS-CoV-2 5' UTR-EcoRI-MluI**

GCGCGCCATTGGGATGGAACTAATACGACTCACTATTATTAAAGGTTTATACCTTCCCAGGTAA
CAAACCAACCAACTTTCGATCTCTTGTAGATCTGTTCTCTAAACGAACTTTAAAATCTGTGTGG
CTGTCACTCGGCTGCATGCTTAGTGCACTCACGCAGTATAATTAATAACTAATTACTGTCGTTG
ACAGGACACGAGTAACTCGTCTATCTTCTGCAGGCTGCTTACGGTTTCGTCCGTGTTGCAGCCG
ATCATCAGCACATCTAGGTTTCGTCCGGGTGTGACCGAAAGGTAAGGAATTCGTTCCATCCCAA
TACGCGT

**MluI-T7-5utr(CoV)-FLUC-3utr(CoV)-HinDIII-AscI (Luciferase fusion construct)**

ACGCGTATTGGGATTAATACGACTCACTATTattaaaggtttataccttcccaggtaacaaacc
aaccaactttcgatctcttgtagatctgttctctaaacgaactttaaaatctgtgtggctgtca
ctcggctgcatgcttagtgcactcacgcagtataattaataactaattactgtcgttgacagga
cacgagtaactcgtctatcttctgcaggctgcttacggtttcgtccgtgttgcagccgatcatc
agcacatctaggtttcgtccgggtgtgaccgaaaggtaag**ATG**GAAGACGCCAAAAACATAAAG

AAAGGCCCGGCGCCATTCTATCCTCTAGAGGATGGAACCGCTGGAGAGCAACTGCATAAGGCTA
TGAAGAGATACGCCCTGGTTCCTGGAACAATTGCTTTTACAGATGCACATATCGAGGTGAACAT
CACGTACGCGGAATACTTCGAAATGTCCGTTCGGTTGGCAGAAGCTATGAAACGATATGGGCTG
AATACAAATCACAGAATCGTCGTATGCAGTGAAAACTCTCTTCAATTCTTTATGCCGGTGTTGG
GCGCGTTATTTATCGGAGTTGCAGTTGCGCCCGCGAACGACATTTATAATGAACGTGAATTGCT
CAACAGTATGAACATTTCGCAGCCTACCGTAGTGTTTGTTTCCAAAAAGGGGTTGCAAAAAATT
TTGAACGTGCAAAAAAAATTACCAATAATCCAGAAAATTATTATCATGGATTCTAAAACGGATT
ACCAGGGATTTCAGTCGATGTACACGTTCGTCACATCTCATCTACCTCCCGGTTTTAATGAATA
CGATTTTGTACCAGAGTCCTTTGATCGTGACAAAACAATTGCACTGATAATGAATTCCTCTGGA
TCTACTGGGTTACCTAAGGGTGTGGCCCTTCCGCATAGAACTGCCTGCGTCAGATTCTCGCATG
CCAGAGATCCTATTTTTGGCAATCAAATCATTCCGGATACTGCGATTTTAAGTGTTGTTCCATT
CCATCACGGTTTTGGAATGTTTACTACACTCGGATATTTGATATGTGGATTTCGAGTCGTCTTA
ATGTATAGATTTGAAGAAGAGCTGTTTTTACGATCCCTTCAGGATTACAAAATTCAAAGTGCGT
TGCTAGTACCAACCCTATTTTCATTCTTCGCCAAAAGCACTCTGATTGACAAATACGATTTATC
TAATTTACACGAAATTGCTTCTGGGGGCGCACCTCTTTCGAAAGAAGTCGGGGAAGCGGTTGCA
AAACGCTTCCATCTTCCAGGGATACGACAAGGATATGGGCTCACTGAGACTACATCAGCTATTC
TGATTACACCCGAGGGGGATGATAAACCGGGCGCGGTCGGTAAAGTTGTTCCATTTTTTGAAGC
GAAGGTTGTGGATCTGGATACCGGGAAAACGCTGGGCGTTAATCAGAGAGGCGAATTATGTGTC
AGAGGACCTATGATTATGTCCGGTTATGTAAACAATCCGGAAGCGACCAACGCCTTGATTGACA
AGGATGGATGGCTACATTCTGGAGACATAGCTTACTGGGACGAAGACGAACACTTCTTCATAGT
TGACCGCTTGAAGTCTTTAATTAAATACAAAGGATATCAGGTGGCCCCCGCTGAATTGGAATCG
ATATTGTTACAACACCCCAACATCTTCGACGCGGGCGTGGCAGGTCTTCCCGACGATGACGCCG
GTGAACTTCCCGCCGCCGTTGTTGTTTTGGAGCACGGAAAGACGATGACGGAAAAAGAGATCGT
GGATTACGTCGCCAGTCAAGTAACAACCGCGAAAAAGTTGCGCGGAGGAGTTGTGTTTGTGGAC
GAAGTACCGAAAGGTCTTACCGGAAAACTCGACGCAAGAAAAATCAGAGAGATCCTCATAAAGG
CCAAGAAGGGCGGAAAGTCCAAACTCGAG**TAG**caatctttaatcagtgtgtaacattagggagg
acttgaaagagccaccacattttcaccgaggccacgcggagtacgatcgagtgtacagtgaaca
atgctagggagagctgcctatatggaagagccctaatgtgtaaaattaattttagtagtgctat
ccccatgtgattttaatagcttcttaggagaatgacaaaaaaaaaaaaaaaaaaaaaaaaaaaa
aaaaAAGCTTATCCCAATGGCGCGCC

**NcoI-His$_6$-PrG-TEV- Met-Ala-(*p*Az)F-eIF4E-XhoI**

CCATGGGTAGCTCACATCATCATCATCATCACTCTTCTGGTCTGGTCCCGCGTGGCTCGCACAT
GCAATACAAACTGATTCTGAACGGTAAAACGCTGAAAGGTGAAACCACGACCGAAGCAGTGGAT
GCGGCCACCGCTGAAAAAGTTTTCAAACAGTACGCCAACGATAATGGCGTGGATGGTGAATGGA
CCTATGATGACGCAACGAAAACCTACACGGTGACCGAAGGTTCCGGCGGTGAAAATCTGTACTT
CCAAGGCCATATGGCG**TAG**ATGGCCACCGTTGAACCGGAAACGACCCCGACGACCAACCCGCCG
CCGGCTGAAGAAGAAAAAACCGAAAGCAACCAGGAAGTCGCGAATCCGGAACATTATATTAAAC
ACCCGCTGCAAAACCGTTGGGCTCTGTGGTTTTTCAAAAACGATAAATCAAAAACGTGGCAGGC
GAACCTGCGCCTGATTTCGAAATTTGATACCGTGGAAGACTTCTGGGCACTGTATAACCACATC
CAACTGAGCTCTAATCTGATGCCGGGTTGCGATTACAGCCTGTTTAAAGACGGCATTGAACCGA
TGTGGGAAGATGAGAAAAACAAACGTGGCGGTCGCTGGCTGATCACGCTGAACAAACAGCAACG
TCGCTCTGATCTGGACCGTTTTTGGCTGGAAACCCTGCTGTGCCTGATTGGCGAAAGTTTCGAT
GACTACTCCGATGACGTTTGTGGTGCGGTGGTTAATGTCCGTGCCAAAGGCGATAAAATTGCAA

TCTGGACGACCGAATGTGAAAACCGCGACGCCGTCACCCATATCGGCCGTGTGTATAAAGAACG
CCTGGGTCTGCCGCCGAAAATTGTTATCGGCTACCAGAGCCACGCAGATACGGCGACCAAATCG
GGCAGCACCACCAAAAATCGTTTCGTTGTGTGACTCGAG

**NcoI-His$_6$-PrG-TEV-Met-Ala-(*p*Az)F-eIF4E(S209D)-XhoI**

CCATGGGTAGCTCACATCATCATCATCATCACTCTTCTGGTCTGGTCCCGCGTGGCTCGCACAT
GCAATACAAACTGATTCTGAACGGTAAAACGCTGAAAGGTGAAACCACGACCGAAGCAGTGGAT
GCGGCCACCGCTGAAAAAGTTTTCAAACAGTACGCCAACGATAATGGCGTGGATGGTGAATGGA
CCTATGATGACGCAACGAAAACCTACACGGTGACCGAAGGTTCCGGCGGTGAAAATCTGTACTT
CCAAGGCCATATGGCG**TAG**ATGCTGGAAATGGCCACCGTTGAACCGGAAACGACCCCGACGCCG
AACCCGCCGACCACCGAAGAAGAAAAAACCGAAAGCAACCAGGAAGTCGCGAATCCGGAACATT
ATATTAAACACCCGCTGCAAAACCGTTGGGCTCTGTGGTTTTTCAAAAACGATAAATCAAAAAC
GTGGCAGGCGAACCTGCGCCTGATTTCGAAATTTGATACCGTGGAAGACTTCTGGGCACTGTAT
AACCACATCCAACTGAGCTCTAATCTGATGCCGGGTTGCGATTACAGCCTGTTTAAAGACGGCA
TTGAACCGATGTGGGAAGATGAGAAAAACAAACGTGGCGGTCGCTGGCTGATCACGCTGAACAA
ACAGCAACGTCGCTCTGATCTGGACCGTTTTTGGCTGGAAACCCTGCTGTGCCTGATTGGCGAA
AGTTTCGATGACTACTCCGATGACGTTTGTGGTGCGGTGGTTAATGTCCGTGCCAAAGGCGATA
AAATTGCAATCTGGACGACCGAATGTGAAAACCGCGAAGCCGTCACCCATATCGGCCGTGTGTA
TAAAGAACGCCTGGGTCTGCCGCCGAAAATTGTTATCGGCTACCAGAGCCACGCAGATACGGCG
ACCAAATCGGGCAGCACCACCAAAAATCGTTTCGTTGTGTGACTCGAG

**NcoI-eIF4G(557–1137)-BamHI** (Codon-optimized for *E. coli*)

CCATGGGGCATCATCACCATCACCACGAGTCAGAAGGGTCGGGAGTCCCGCCAAGACCCGAAGA
GGCGGATGAGACATGGGATTCGAAAGAGGATAAAATCCATAATGCTGAAAATATACAACCTGGA
GAGCAGAAATACGAGTATAAATCGGACCAATGGAAACCATTGAACTTAGAGGAAAAAAAGCGCT
ACGATCGTGAATTCCTTTTGGGGTTTCAATTCATCTTTGCATCGATGCAAAAGCCAGAGGGCCT
GCCACATATCAGTGATGTTGTACTTGACAAGGCGAATAAGACACCGTTAAGACCCTTAGATCCT
ACGCGTTTACAAGGAATAAATTGCGGACCAGATTTTACGCCTAGCTTCGCAAATCTTGGTAGAA
CAACCCTGAGCACACGCGGCCCTCCCAGAGGTGGCCCAGGGGGGGAGCTGCCACGCGGACCAGC
AGGTTTAGGGCCAAGACGTTCACAACAGGGGCCGCGGAAAGAACCTCGCAAAATCATAGCGACG
GTATTGATGACAGAGGATATTAAATTGAATAAAGCAGAAAAGGCGTGGAAGCCATCTTCGAAAC
GCACAGCCGCTGATAAGGATCGGGGTGAGGAGGACGCCGATGGCAGCAAAACACAAGATCTTTT
CCGGCGGGTCCGTAGCATATTAAACAAGCTTACTCCACAGATGTTCCAACAGCTGATGAAGCAA
GTGACACAGTTGGCGATTGACACAGAAGAACGGCTTAAGGGGGTCATTGATTTGATATTTGAAA
AAGCCATTAGTGAGCCCAACTTCAGCGTAGCTTACGCTAATATGTGTCGTTGTTTAATGGCGCT
TAAAGTTCCGACTACGGAGAAACCGACCGTAACAGTGAATTTCCGTAAATTGCTTTTAAACCGT
TGTCAAAAGGAATTCGAAAAAGACAAGGATGACGACGAGGTTTTCGAGAAAAAACAAAAAGAGA
TGGACGAGGCCGCTACAGCAGAGGAAAGAGGTAGATTAAAGGAAGAATTAGAAGAGGCCCGCGA
CATCGCGCGGCGGAGAAGCTTGGGAAATATCAAATTTATAGGGGAACTTTTCAAATTGAAAATG
CTGACTGAGGCAATCATGCATGACTGCGTTGTTAAGCTGTTAAAGAACCATGACGAGGAGTCGT
TAGAGTGCTTGTGCCGGTTACTTACAACAATTGGCAAAGATCTGGACTTTGAAAAGGCGAAGCC
ACGCATGGACCAATACTTTAACCAGATGGAAAAAATTATAAAGGAAAAAAAAACCTCGAGCCGG

ATAAGATTTATGTTGCAAGACGTACTGGATTTACGTGGGTCTAACTGGGTTCCCCGTCGGGGGG
ACCAGGGTCCGAAGACCATAGATCAAATCCACAAGGAAGCGGAAATGGAAGAGCATCGTGAACA
TATAAAGGTTCAACAATTAATGGCCAAAGGGTCAGACAAGCGGCGTGGCGGTCCCCCTGGGCCG
CCCATCTCAAGAGGATTACCATTGGTTGACGACGGCGGATGGAATACCGTACCTATTAGCAAGG
GCAGCCGCCCTATAGACACATCGCGGCTTACTAAGATTACGAAACCAGGGTCGATTGATTCCAA
CAACCAACTGTTTGCACCAGGCGGTCGCTTGAGTTGGGGCAAGGGTTCATCGGGCGGTTCGGGA
GCAAAGCCCTCTGATGCCGCCTCGGAAGCAGCGAGACCAGCAACGTCTACTCTTAACCGGTTTT
CTGCTCTGCAACAAGCGGTCCCTACCGAGTCTACGGATAACTAGGATCC
